# Reduced precipitation alters microbial availability and uptake of grassland rhizosphere carbon

**DOI:** 10.64898/2026.04.10.717538

**Authors:** Nayela Zeba, Katerina Y. Estera-Molina, Siyang Jian, Jing Yan, Steven J. Blazewicz, Aaron Chew, Nhu H. Nguyen, Jizhong Zhou, Jennifer Pett-Ridge, Mary K. Firestone, Mengting Yuan

**Affiliations:** Pacific Biosciences Research Center, University of Hawaiʻi at Mānoa, Honolulu, HI 96822 USA; Department of Environmental Science, Policy, and Management, University of California, Berkeley, CA 94720 USA; Physical and Life Sciences Directorate, Lawrence Livermore National Laboratory, Livermore, CA 94551 USA; Institute for Environmental Genomics, University of Oklahoma, Norman, OK 73019 USA; Department of Tropical Plant and Soil Sciences, University of Hawaiʻi at Mānoa, Honolulu, HI 96822 USA; School of Civil Engineering and Environmental Sciences, University of Oklahoma, Norman, OK 73019 USA; School of Computer Sciences, University of Oklahoma, Norman, OK 73019 USA; Earth and Environmental Sciences, Lawrence Berkeley National Laboratory, Berkeley, CA 94720 USA; Life & Environmental Sciences Department, University of California Merced, Merced, CA 95343 USA; Innovative Genomics Institute, University of California Berkeley, Berkeley, CA 94720 USA

**Keywords:** rhizodeposit C, rhizosphere microbiome, precipitation reduction, microbial co-occurrence networks, Mediterranean grassland

## Abstract

Climate change intensifies grassland drying, which is predicted to decrease soil organic carbon (SOC) storage. Yet how reduced soil water affects the transport and processing of carbon in the rhizosphere — a major source of SOC supporting microbial activities — remains poorly resolved under field conditions. We traced living root-derived carbon (rhizodeposit C) into soils and microbes using ¹³CO₂ labeling in a multi-year precipitation manipulation experiment in a California Mediterranean annual grassland dominated by *Avena barbata*. Under halved precipitation, season-averaged soil water content was 20% lower and soil matric potential more than threefold lower, without inducing detectable signs of plant stress. Quantitative stable isotope probing (qSIP) of rhizosphere microbial DNA revealed that up to 68% more bacterial and 13% more fungal taxa incorporated rhizodeposit C, and that microbial uptake of rhizodeposit C increased more over time under reduced precipitation, even as total microbial biomass and community composition were unchanged. ¹³C-informed co-occurrence networks of these consumers were larger and more connected under reduced precipitation. The surrounding soils contained less ^13^C, and modeled solute transport by soil water was 32-40% lower, implying that rhizodeposit C was retained and intensively processed nearer to roots rather than transported to the wider soil matrix. Our evidence supports that reduced precipitation concentrates rhizodeposit C near roots, increasing the number of distinct consumers and intensifying associations among them. This represents an overlooked mechanism linking precipitation reduction to rhizodeposit C cycling. These findings show that precipitation reduction may reshape rhizosphere C processing before apparent plant stress response, with potential consequences for the fate of rhizodeposit C in grassland soils.

## 1. Introduction

Grasslands cover 23% of the terrestrial land area (MacDougall et al., 2026) and store between 10–30% of global soil organic carbon (SOC) (Scurlock and Hall, 1998; Conant et al., 2017) but face intensifying challenges from climate change through reduced and more erratic precipitation. These shifts may have significant implications for the persistence of SOC (Arca et al., 2021; Deng et al., 2021). SOC stocks reflect the balance between plant-derived C inputs from root exudates and litter, and microbial decomposition; both of which decrease under reduced precipitation and have been linked to SOC destabilization and net ecosystem C loss (Deng et al., 2021; Zhang and Xi, 2021; Bai and Cotrufo, 2022). While the influence of soil water on litter decomposition is well documented (Sanaullah et al., 2012; Allison et al., 2013; Zhao et al., 2025), how altered precipitation influences SOC dynamics via C flow through the rhizosphere, the soil surrounding living roots, remains less clear.

Grasses allocate about one-third of photosynthetic C belowground before senescence, with a substantial portion entering soil in rhizodeposit, organic molecules released from living roots (Pausch and Kuzyakov, 2018). Rhizodeposit fuels microbial activities that regulate SOC formation, stabilization, and mineralization (Paterson et al., 2007; Villarino et al., 2021; Teixeira et al., 2024). A key component of the stabilization process is mineral-associated organic matter (MAOM), the largest and slowest-cycling SOC pool (Lehmann et al., 2020) that forms in part through association of microbial processed rhizodeposit C with soil minerals (Fossum et al., 2022; Sokol et al., 2024). Rhizodeposit C is processed rapidly: fresh root exudates are transformed within days to weeks into microbial metabolites and necromass (Balasooriya et al., 2014; Huang et al., 2020) before entering higher trophic levels of soil food web or associating with minerals to form MAOM. Yet field-based studies that track the flow and microbial uptake of rhizodeposit C under realistic soil water dynamics remain scarce, limiting predictions of SOC responses to changing precipitation regimes.

Soil water influences rhizodeposit C processing through mechanisms operating at multiple scales. For plants, reduced water availability affects growth, root activity, and rhizodeposition. Grasses start to become stressed when the matric potential drops below −0.7 MPa (Fu et al., 2022). Stressed plants may reduce total C inputs to soils, even as the fraction of fixed C allocated to root exudates increases or remains unchanged (Preece and Peñuelas, 2016; Karlowsky et al., 2018; Wang et al., 2021b). For microbes, especially bacteria, movement through soil becomes restricted under moderate water limitation (around −0.3 MPa) (Gentry et al., 2021), and as water availability decreases further, physiological activity and enzyme function are constrained (Gomez et al., 2020), limiting C consumption. Beyond these biological effects, physical transport of dissolved rhizodeposit C through soil slows as soils dry (Wang et al., 2021a), potentially constraining MAOM formation (Manzoni et al., 2012). This drying also fragments continuous water films into disconnected microsites, altering how microbial communities access and compete for substrates (Treves et al., 2003; Tecon and Or, 2017; Tecon et al., 2018).

Species of the genus *Avena* (common oatgrass) provide an excellent system for studying rhizosphere C cycling because of their widespread global distribution across temperate grasslands, their substantial contribution to ecosystem productivity, and their close relationship to agronomically important crops (Lu et al., 2025). In *Avena* rhizosphere, bacteria and fungi play central, complementary roles in consuming and transforming rhizodeposit C, with bacteria specializing in fresh exudate degradation and fungi in breaking down complex plant polymers, and fungal biomass itself serving as a C substrate for bacterial consumers (Shi et al., 2015; Zhalnina et al., 2018; Nuccio et al., 2021; Pett-Ridge et al., 2021; Yuan et al., 2021; Kakouridis et al., 2024). Microbes from different functional guilds share or partition niche spaces, cooperate or compete in complex networks to influence C processing, relocation, and transformation around roots (Shi et al., 2016; Nuccio et al., 2020; Pett-Ridge et al., 2021; Yuan et al., 2021). Co-occurrence networks are widely used to study microbial associations across environmental gradients (Barberán et al., 2012). Water reduction has been shown to restructure rhizosphere ecological networks with stronger effects on bacteria than on fungi or protists, although changes in network complexity and stability vary across systems (de Vries et al., 2018; J. Wang et al., 2023). Correlation-based networks built from relative abundances may reflect shared environmental responses or indirect associations rather than direct interactions (Barberán et al., 2012; Faust et al., 2015; Hirano and Takemoto, 2019; Guseva et al., 2022). ^13^C quantitative stable isotope probing (qSIP) is a complementary approach that directly links taxa to substrate assimilation. ^13^CO_2_ can be used to track freshly plant-fixed C into microbial DNA to quantify microbial consumption of plant-derived C (Hungate et al., 2015; Nuccio et al., 2022), enabling mapping of relevant taxa in microbial networks (Slanzon et al., 2025).

Our study aims to evaluate how reduced precipitation alters grassland rhizodeposit C processing by its microbial consumers. We answer this question with whole-plant field-based ^13^CO_2_ labeling followed by rhizodeposit ^13^C tracing into microbial communities and surrounding soils under contrasting precipitation levels. We conducted this work in a California annual grassland, representative of Mediterranean-climate grasslands that occupy ∼4% of global grassland area (Dixon et al., 2014), where frequent extreme dry-to-wet precipitation swings are projected to intensify growing-season water stress (Swain et al., 2018; Felton et al., 2020). We labeled *Avena*-dominated vegetation with ¹³CO₂ for eight days where precipitation received was either the 50-year average or half that amount for three years. Using qSIP, we quantified bacterial and fungal incorporation of rhizodeposit ^13^C immediately after labeling (March 12, 2020; exponential plant growth) and four weeks later (April 13, 2020; peak plant biomass), distinguishing early from later-stage consumers. We constructed bacteria-fungi co-occurrence networks for rhizodeposit C consumers. Finally, we measured ¹³C distribution across plant and soil pools, together with solute-transport modeling, to interpret the movement of rhizodeposit C through soil and the physical context of these microbial responses.

## 2. Materials and Methods

### 2.1. Precipitation manipulation field experiment

This study was conducted at the Hopland Research and Extension Center (HREC) in Mendocino County, California, USA (39.004 °N, –123.086 °W; elevation ≈ 210 m), as part of a multi-year precipitation manipulation experiment established in fall 2017 (Fossum et al., 2022). The site has a Mediterranean climate with hot, dry summers (mean July–September temperature ≅ 21 °C, mean maximum ≅ 33 °C) and mild, rainy winters, receiving about 75% of its mean annual precipitation between November and February. Vegetation is dominated by the annual wild oat *Avena barbata*, with scattered grasses (*Bromus, Festuca*) and forbs (*Plantago, Trifolium*). Soils belong to the Squawrock-Witherell complex: dominantly gravelly loam (Squawrock) with loam (Witherell) on 15-50% west-facing slopes (USDA-NRCS).

In fall 2017, sixteen 1.8 × 1.8 m plots were established and lined to a depth of 1 m with pond liner to minimize lateral water movement between plots and the surrounding soil. Thereafter, eight randomly assigned plots received combined natural precipitation, spring water irrigation and rain shelter treatment to simulate the 50-year historical average precipitation. The other eight received half that amount, by a watering management schedule that targeted precipitation levels during the plant growing season on a bi-weekly basis. Soil volumetric water content (*θ_v_*) at 0-5 cm depth was recorded every 2 hours using EC-5 soil moisture probes connected to EM5b data loggers (Decagon Devices, Pullman, WA). Maximum plant height was estimated ten times throughout the growing season by measuring the height of three tallest individual *Avena* plants in each plot.

### 2.2. ¹³CO₂ pulse labeling

Data presented here were collected from an 8-day ^13^CO_2_ labeling campaign conducted in spring 2020, following 2.5 years of precipitation manipulation. Each plot was equipped with two PVC collars fitted with Plexiglas dividers, inserted 15 cm into the soil, to create two semicircular wedges for destructive sampling at two time points (Fossum et al., 2022). Cylindrical labeling chambers were mounted on the collars and supplied with either ¹²CO₂ or 99 atom% ¹³CO₂ (Cambridge Isotope Laboratories, Tewksbury, MA) for the entire daytime period when photosynthetically active radiation (PAR) exceeded 50 µmol m⁻² s⁻¹ over the labeling period (March 3-11, 2020), targeting the exponential growth stage of *Avena* grasses. CO_2_ delivery and headspace concentrations in the chambers were monitored and controlled in real time with the Dynamic Ecosystem Labeling (DEL) system (Fossum et al., 2022). Headspace CO_2_ concentrations were maintained between 400-700 ppm during the day. A total of twenty-two chambers were deployed (11 ¹³CO₂, 11 ¹²CO₂), comprising five ¹²C/¹³C pairs for the average precipitation treatment and six pairs for the reduced precipitation treatment.

### 2.3. Sample collection and measurement

Shoots, roots, and soil were harvested at two timepoints: March 12, 2020 (one day post-labeling period) and April 13, 2020 (four weeks post-labeling). At each time point, one semicircular wedge (0-15 cm depth) from each collar was destructively sampled using ethanol and Eliminase-sterilized tools. Root samples with attached soil were carefully separated, immediately placed on dry ice, and stored at −80 °C until processing. Soil tightly adhering to these roots was separated using a standardized root-washing and extraction protocol (Shi et al., 2015), and designated as “rhizosphere”. Briefly, approximately 2 g of root material was transferred into 50-mL conical tubes containing 6 mL of Lifeguard Soil Preservation Solution (Qiagen, Germantown, MD) on wet ice. The tubes were vortexed horizontally for 1 minute at maximum speed, followed by centrifugation at 4,000 rpm for 15 minutes at 4 °C. Washed roots were manually removed using sterile inoculation loops and tweezers, transferred to 15-mL tubes. The remaining soil suspension was filtered through a sterile 1-mm flow mesh (Thomas Scientific, Swedesboro, NJ) into pre-weighed 50-mL tubes to remove remaining root debris. The soil fraction was pelleted via centrifugation for 5 minutes, the supernatant was discarded, and the resulting rhizosphere soil pellet was flash-frozen on dry ice and stored at −80 °C until PLFA analysis and DNA extraction. A subsample of the surrounding soil that had been picked free of major roots was stored separately and categorized as “surrounding soil”. The surrounding soil was different from “bulk soil”, since this grassland is densely rooted during the growing season and all surface soil had likely been under the influence of roots in the field.

At both harvest timepoints, aboveground biomass was clipped, dried at 65 °C to constant weight, and weighed. Washed roots were also dried. Both dried root and shoot biomass were ground to a fine powder and encapsulated in tins for δ¹³C analysis. Roots were processed only for δ¹³C analysis, not for biomass quantification, because fine-root recovery during sample collection was too variable for reliable root biomass estimates. Soil gravimetric water content (*θ_g_*) was measured on homogenized surrounding soil samples by drying at 65 °C to constant weight. The dried surrounding soils were then ground for δ¹³C analysis as above. All δ¹³C measurements were performed on an elemental analyzer interfaced with an isotope ratio mass spectrometer (EA-IRMS) at the UC Davis Stable Isotope Facility (Davis, CA) and the Stable Isotope Measurement Facility, University of Oklahoma (Norman, OK).

### 2.4. DNA qSIP analysis

To identify ¹³C-rhizodeposit bacterial and fungal consumers, DNA was extracted from two 0.5 g aliquots of rhizosphere soil per sample using a modified phenol-chloroform co-extraction protocol (Griffiths et al., 2000; Shi et al., 2015) and processed using the HT-SIP pipeline (Nuccio et al., 2022) for density fractionation, amplicon sequencing, and qSIP.

*DNA extraction and density fractionation:* Briefly, samples were homogenized in Lysing Matrix E tubes (MP Biomedicals, Solon, OH) with CTAB extraction buffer, followed by phenol-chloroform purification, PEG precipitation, ethanol washing, and lithium chloride separation from RNA. DNA pellets were air-dried, resuspended in molecular-grade water, and quantified for purity using a NanoDrop One spectrophotometer (Thermo Fisher Scientific, Waltham, MA) and for concentration using a Qubit 3.0 fluorometer (Thermo Fisher Scientific, Waltham, MA). For each sample, 1-5 µg of DNA was loaded into a cesium chloride density gradient and ultracentrifuged at Lawrence Livermore National Laboratory to separate DNA by buoyant density. Each gradient yielded 22 fractions, which were combined into four final fractions per sample based on density to ensure sufficient DNA yield for both bacterial and fungal amplicon library preparation. The density of each combined fraction was calculated as the proportional average of its constituent fractions.

*Amplicon sequencing:* For each sample, we sequenced the four density-resolved SIP fractions for qSIP analysis and the unfractionated DNA for total community profiling. The V4 region of the bacterial and archaeal 16S rRNA gene was amplified using the 515F/806R primer pair (Caporaso et al., 2011), and the ITS2 region of the fungal rRNA gene was amplified using the ITS5.8S-Fun/ ITS4-Fun primer pair (Taylor et al., 2016). Amplicons were sequenced on an Illumina MiSeq platform (Illumina, San Diego, CA) with paired-end 2×250 bp reads for 16S and 2×300 bp reads for ITS2 across three sequencing runs at the Institute for Environmental Genomics, University of Oklahoma (Norman, OK).

*Bioinformatic processing:* Paired-end 16S and ITS2 reads from the three Illumina MiSeq runs were processed using QIIME 2 v2024.5 (Bolyen et al., 2019) on the Savio high-performance computing cluster (University of California, Berkeley, CA). Runs were processed separately: FASTQ files were imported, demultiplexed, and primer-trimmed using the q2-cutadapt plugin (Martin, 2011). Quality filtering, chimera removal, and amplicon sequence variant (ASV) inference were performed with the DADA2 plugin (Callahan et al., 2016). Pre-filtering read counts ranged from 36,676-178,281 for 16S and 30,880-212,575 for ITS2. Feature tables and representative sequences were merged across runs (feature-table merge with method = sum, and feature-table merge-seqs), and ASVs observed fewer than two times in fewer than two samples were removed. Taxonomy for 16S ASVs was assigned using classify-sklearn with the pre-trained SILVA 138 SSURef NR99 OTU classifier (Bokulich et al., 2018; Kaehler et al., 2019); chloroplast, mitochondrial, and archaeal sequences were excluded, yielding 7,015 unique bacterial ASVs. ITS2 ASVs were classified with the same method using a classifier trained on the UNITE v10.0 database (release 04.04.2024), with dynamic clustering (97-99% identity) and exclusion of global singletons (Abarenkov et al., 2024), yielding 4,018 unique fungal ASVs. Post-filtering read counts ranged from 13,493-89,715 (16S) and 42,691-105,932 (ITS2) per unfractionated sample, and from 12,025-70,956 (16S) and 8,860-155,023 (ITS2) per SIP-fractionated sample (Tables S1–S4).

*qSIP analysis:* The qSIP analysis identifies ASVs that have incorporated ¹³C and quantifies their incorporation by converting taxon-specific shifts in DNA buoyant density into ¹³C excess atom fraction (EAF) (Hungate et al., 2015). Analyses were performed using the *qSIP2* R package (https://github.com/jeffkimbrel/qSIP2). Within each precipitation × timepoint group, ASVs were retained if detected in at least four biological replicates and at least three density fractions per replicate in both ¹³C-labeled samples and their ^12^C counterparts. This filtering retained 4-7% of 16S ASVs (69-97% of reads; Table S3) and 1-3% of ITS2 ASVs (83-96% of reads; Table S4). Bootstrap resampling with 1,000 iterations was used to calculate 90% confidence intervals (CIs) for mean EAF values. ASVs with a mean EAF > 0 (66-94% of 16S ASVs and 70-86% of ITS2 ASVs) were considered ^13^C-enriched and categorized as ¹³C-rhizodeposit consumers.

We applied the mean EAF > 0 throughout all analyses because it captures both strong and weak ^13^C incorporators, reducing selection bias when estimating continuous community-level metrics. Although the more conservative criterion of lower CI > 0 generated similar contrasts of the numbers and enrichment levels of ¹³C-rhizodeposit consumers across treatments (Fig. S4), it would preferentially retain only strongly enriched ASVs in EAF-weighted relative abundances used in network construction (described in section 2.6) and bias network topology.

To calculate the effect of reduced precipitation on rhizodeposit-derived C incorporation over time, we focused on persistent consumers (i.e., ASVs with mean EAF > 0 in both water treatments at both timepoints). For each persistent consumer, we calculated the relative effect of reduced precipitation on its ^13^C enrichment (*R_EAF_*) at each timepoint as:

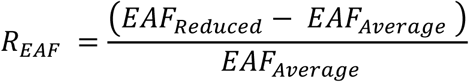

Similarly, we calculated the relative effect of reduced precipitation on relative abundance (*R_RA_*) for each persistent consumer at each timepoint as:

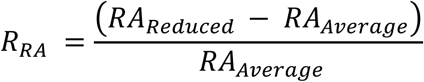

*R_EAF_* > 0 indicates higher ¹³C EAF under reduced compared with average precipitation at that timepoint; *R_RA_* > 0 indicates higher relative abundance under reduced compared with average precipitation at that timepoint

To visualize temporal dynamics, we plotted *R_EAF_* and *R_RA_* for each persistent consumer at both timepoints, using arrows to indicate the magnitude and direction of the shift from the first to the second timepoint.

### 2.5. PLFA Analysis

The PLFA analysis was performed in the Institute for Environmental Genomics at the University of Oklahoma (Norman, OK). Lipids were extracted from the rhizosphere following a modified Bligh–Dyer method (Buyer and Sasser, 2012). Briefly, soil samples were vacuum-dried (SPD1010 SpeedVac System, Thermo Scientific, MA) and sieved to remove rocks and large debris. One gram of dried soil was sonicated in a Bligh–Dyer extractant consisting of methanol, chloroform, and 50 mM K₂HPO₄ buffer (2:1:0.8, v/v/v). An internal standard of 19:0 phosphatidylcholine (Avanti Polar Lipids, AL) dissolved in a 1:1 chloroform:methanol solution (40.9 mg per 20 mL) was added at a rate of 0.5 μL per mL of extractant immediately prior to use. The chloroform phases were collected, and phospholipids were separated from neutral lipids and glycolipids using a 96-well solid-phase extraction plate (Sigma-Aldrich, MO). The phospholipid fraction was trans-esterified to fatty acid methyl esters (FAMEs) and analyzed using an Agilent 6890N gas chromatograph (Agilent Technologies, DE). Peaks were identified with the Sherlock PLFA Analysis software 6.3 (Biolog Inc., CA) using PLFAD1 calibration standards.

Peaks were quantified as pmol of fatty acid per gram of dry soil, normalized to the 19:0 internal standard. Fatty acids were assigned to microbial groups by the Sherlock PLFA Analysis software: gram-negative bacteria (monounsaturated and cyclopropyl 17:0 and 19:0 FAs), gram-positive bacteria (iso- and anteiso-branched FAs), actinomycetes (10-methyl FAs), anaerobic bacteria (dimethyl acetal FAs), common fungi (18:2ω6c), and arbuscular mycorrhizal fungi (AMF) (16:1ω5c). Total bacterial biomass was calculated as the sum of gram-negative, gram-positive, actinomycete, and anaerobic bacterial biomarkers, and total fungal biomass was calculated as the sum of fungal and AMF biomarkers.

### 2.6. Network analysis

Microbial co-occurrence networks were constructed for each precipitation × timepoint combination using unfractionated DNA to identify potential ecological associations within bacterial and fungal communities. We used SPIEC-EASI to infer conditional dependence networks, an approach well-suited to amplicon sequencing because it accommodates compositional and sparse data structures (Kurtz et al., 2015). Two network types were constructed per combination group: (1) ^13^C-rhizodeposit community networks (“^13^C networks”), built using EAF-weighted relative abundances (RA) of qSIP-identified consumers to capture associations among bacteria and fungi actively involved in rhizodeposit C consumption; and (2) total community networks based solely on relative abundances.

*Pre-filtering of ASVs for network construction:* For ^13^C networks, ASVs in unfractionated communities were retained if they were ^13^C-enriched (i.e., mean EAF > 0 from qSIP). The per-sample relative abundances of these ASVs were weighted by mean EAF values. This produced an EAF-weighted RA table in which each value represents an ASV’s relative importance in rhizodeposit C processing; i.e., higher values indicate ASVs that are both abundant and highly ¹³C-enriched. Separate 16S and ITS2 EAF-weighted RA tables were merged and re-normalized so that each sample summed to one across both bacterial and fungal taxa, preserving compositionality prior to network construction. EAF filtering retained 5-9% of 16S ASVs and 1.7-2% of ITS2 ASVs, capturing 39-52% and 40-49% of total 16S and ITS reads respectively (Table S6-S7). For total community networks, ASVs were retained only if present in at least 80% of samples within each combination group, ensuring reproducibility and robust estimation of ecological associations. This prevalence filter retained 6-10% of 16S ASVs and 3.6-5.8% of ITS2 ASVs, accounting for 42-54% and 53-62% of total 16S and ITS reads respectively (Table S9-S10).

*Network construction and topology:* All networks were constructed using the Meinshausen-Bühlmann (MB) neighborhood selection method in the *SpiecEasi* (Kurtz et al., 2024) R package. Edge signs were extracted via the maxabs symmetrization method. The algorithm tested a regularization path of 50 λ values for ¹³C networks and 100 λ values for total community networks (λ_min_/λ_max_ = 0.01). Optimal λ selection was performed independently for each treatment × timepoint combination using the Stability Approach to Regularization Selection (StARS) with 100 subsampling replicates and a stability threshold of 0.05. This calibrated network sparsity separately for each dataset, ensuring that topological comparisons reflect biological rather than regularization-driven differences. Network topology was evaluated using the *igraph* R package (Csárdi et al., 2025). Global topological metrics included the number of nodes and edges, edge sign distribution (positive vs. negative), average node degree, global clustering coefficient, and modularity calculated via the fast-greedy community detection algorithm with absolute edge weights to satisfy the requirement for non-negative weights.

*Keystone species identification:* Keystone taxa were identified using 90th percentile thresholds of degree and betweenness centrality to define three functional roles: hubs (degree ≥ 90th percentile), brokers (betweenness ≥ 90th percentile), and broker-hubs (meeting both criteria). Betweenness was calculated using unweighted shortest paths. Bacterial-fungal connectors were defined as nodes with degree ≥ 5 in which more than 50% of connections linked bacteria to fungi. Taxa meeting criteria for both broker-hub and bacterial-fungal connector were classified primarily as broker-hubs.

### 2.7. Soil hydraulic properties and rhizodeposit C transport modeling

To explore whether restricted solute transport could contribute to the lower ¹³C we observed in surrounding soil under reduced precipitation, we estimated soil water flow and transport parameters from intact soil cores collected adjacent to experimental plots (n = 3) and used these parameters and measured soil water content to model relative solute transport under reduced vs. average precipitation treatments. Each core was equilibrated on a pressure-plate apparatus at six matric potentials (*ψ_m_*, MPa). At each step, gravimetric water content was converted to volumetric water content (*θ_v_*, cm³ cm⁻³) using replicate-specific bulk density, yielding one *θ_v_*-*ψ_m_* curve per core.

Each set of measurements was fit to the van Genuchten model (Van Genuchten, 1980) by minimizing the root-mean-square error between measured and predicted *θ_v_*:

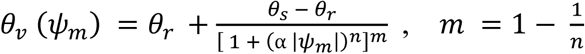

where *Θ_s_* is *θ_v_* at saturation (equal to total porosity calculated from bulk density), *Θ_r_* is residual water content, and ⍺ (MPa⁻¹), *n*, and *m* are empirically fitted parameters optimized for each replicate.

Relative hydraulic conductivity *K_r_* was computed using the Mualem-van Genuchten function:

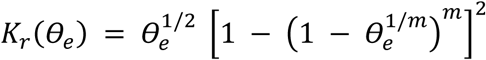

where *K_r_* is dimensionless relative conductivity and *Θ_e_* is effective saturation defined by

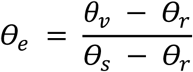

Prior to analysis, *θ_v_* readings exceeding the maximum *Θ_s_* (0.459 cm³ cm⁻³) were removed as sensor artifacts (1.56 % of total records). Data were aggregated to daily means per plot. Each daily *θ_v_* record was then combined with three replicate sets of fitted van Genuchten parameters to produce parallel estimates of *ψ_m_* and *K_r_*. These estimates were averaged across parameter sets to obtain a single daily mean per plot for subsequent analyses.

Using these *θ_v_* records, we estimated three dominant transport parameters governing solute transport: effective molecular diffusion (*D_l_*), pore-water velocity (*V*), and hydrodynamic dispersion (*D*_ℎ_). Calculations followed the framework of Hillel (Hillel, 1975) and were performed for the full growing season and for the period between start of labeling and last sampling (March 3-April 13, 2020). The assumptions and limitations for this simplified model are discussed in section 4.2.

Effective molecular diffusion was calculated following Millington and Quirk (Millington and Quirk, 1961):

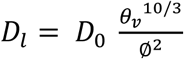

where *D*_0_ is the free-water molecular diffusion coefficient (assumed 0.864 cm² day⁻¹, representative of small organic solutes at 25 °C), and ∅ is total porosity derived from bulk density.

Pore-water velocity and hydrodynamic dispersion were derived from the bulk water flux (*J_w_*) using the Darcy-Buckingham equation:

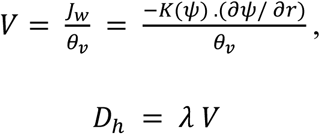

where *K*(*ψ*) is the unsaturated hydraulic conductivity (calculated as *K_s_* ⋅ *K_r_*), *σψ*/ *σr* is the hydraulic gradient, and *λ* is dispersivity (assumed 2 cm at the plot scale (Vanderborght et al., 2000)). Saturated hydraulic conductivity (*K_s_*) was assumed to be uniform across replicates, and a constant unit hydraulic gradient (*σψ*/ *σr* = 1) was assumed to compare relative transport potential between precipitation treatments.

### 2.8. Statistical analyses

All statistical tests were performed in R v4.3.2(R Core Team, 2024).

Soil volumetric water content (*θ_v_*) and matric potential (ψ_m_) were modeled as continuous time-series with linear mixed-effects models including precipitation treatment, time (days since the start of the growing season, scaled), and their interaction as fixed effects; plot as a random intercept; and an AR(1) correlation structure to account for temporal autocorrelation in repeated measures (*nlme* package (J Pinheiro et al., 2023)). Interaction terms were evaluated using likelihood-ratio tests (LRT).

Soil transport parameters (molecular diffusion, average pore-water velocity, and hydrodynamic dispersion) were compared between treatments using Welch’s t-tests on plot-level means over the full growing season and the 42-day labeling-to-last sampling period (March 3-April 13, 2020), with pore-water velocity and hydrodynamic dispersion were log-transformed prior to analysis.

Gravimetric water content (*θ_g_*), atom % ¹³C in surrounding soil, atom % ^13^C in plant tissue, plant biomass, grass height, and bacterial and fungal PLFA biomass were each analyzed using two-way mixed-effects ANOVA with precipitation, timepoint, and their interaction as fixed effects and plot as a random intercept (*lmerTest* package (Kuznetsova et al., 2017)), using Satterthwaite’s method for denominator degrees of freedom. When the interaction term was non-significant (p > 0.05), main effects were examined using estimated marginal means (*emmeans* package (Lenth, 2025)). For grass height, which was measured at ten timepoints during the growing season, date was treated as a categorical factor to enable treatment comparisons at specific growth stages via post hoc tests.

Across all mixed-effects analyses, response variables were log-transformed when needed to meet model assumptions, with normality of residuals assessed by Shapiro-Wilk and homogeneity of variance by Levene’s tests. Treatment means are reported on the original (untransformed) scale.

Community composition of rhizodeposit C-consuming bacteria and fungi was compared between precipitation treatments and timepoints using PERMANOVA on Bray-Curtis dissimilarities calculated from relative abundance data (*phyloseq* (McMurdie and Holmes, 2013) and *vegan* (Oksanen et al., 2022) packages). To restrict analysis to consumers, we subset the unfractionated DNA relative abundance dataset to include only ASVs identified as ^13^C-enriched (mean EAF > 0 from qSIP). The precipitation effect was tested at the plot level by aggregating samples within each plot into a single centroid (one centroid per plot) and comparing dissimilarities between precipitation treatments. The temporal effect was tested across all samples with permutations restricted within plots (strata = plot) to account for repeated measures. Community differences were visualized using principal coordinates analysis (PCoA) of Bray-Curtis dissimilarities, and homogeneity of multivariate dispersions was confirmed using betadisper (*vegan*).

## 3. Results

### 3.1. Soil water availability and plant response

Halved precipitation reduced soil water availability throughout the growing season (∼November through May; Fig. 1A, B); season-averaged soil volumetric water content (θ_v_) was 20% lower (*t*_14_ = −3.83, *p* = 0.002) and modeled soil matric potential (ψ_m_) was more than threefold lower (−369 vs. −1164 kPa; *t*_14_ = 3.90, *p* = 0.002), with more pronounced differences as the growing season progressed. By mid-April, gravimetric water content (θ_g_) was 42% lower in reduced precipitation soils (precipitation × time: *F*_1, 17.8_ = 28.23, *p* < 0.001; Fig. 1C).

**Fig. 1.**
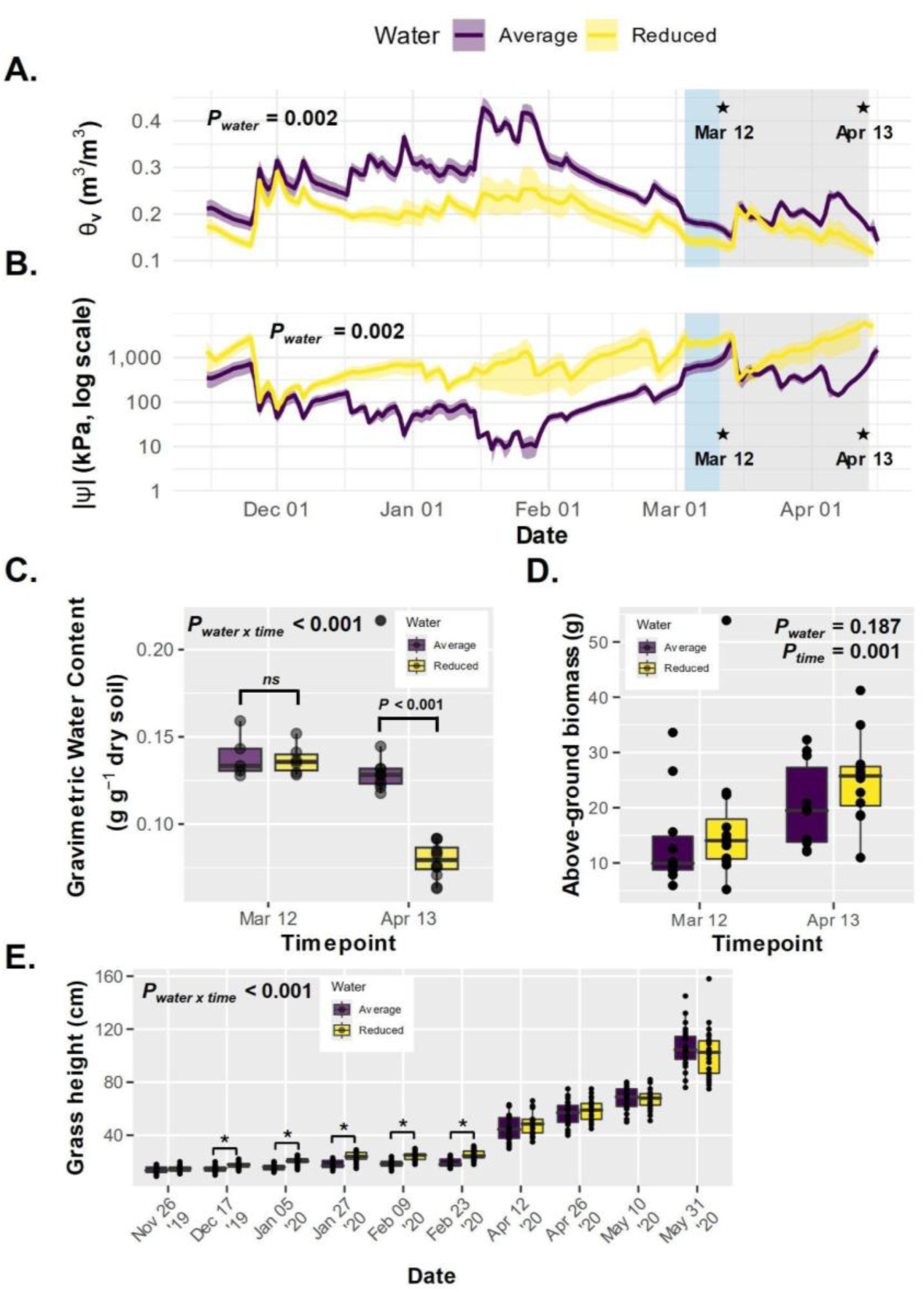
Soil water availability and plant response under contrasting precipitation treatments in a California annual grassland. **(A)** Volumetric water content (θ_v_) and **(B)** modeled matric potential (|ψ_m_|, log scale) from November 2019-April 2020, shown as daily means ± SE for average (purple, *n* = 8) and reduced (yellow, *n* = 8) precipitation plots. Blue shading marks the labeling period (March 3-11); gray shading marks the post-labeling period through final sampling (March 11-April 13); stars indicate sampling timepoints. **(C)** Gravimetric water content (θ_g_) under average (purple, *n* = 5 for March 12 and *n* = 10 for April 13) and reduced (yellow, *n* = 6 for March 12 and *n* = 12 for April 13) precipitation, and **(D)** Dry above-ground biomass under average (purple, *n* = 10) and reduced (yellow, *n* = 12) precipitation at two timepoints (March 12 and April 13, 2020). **(E)** Maximum grass height measured on the three tallest plants within each plot across the 2019-2020 growing season (average, purple, *n* = 24; reduced, yellow, *n* = 24); asterisks indicate dates at which precipitation treatments differed significantly (p < 0.05). Statistical results are from linear mixed-effects models with AR(1) correlation structure (panels A and B), two-way mixed-effects ANOVA (panels C and D), and a linear mixed-effects model with pairwise treatment comparisons at each date (panel E).

Plants showed no visible signs of physiological stress under reduced precipitation: shoot biomass was on average 23% higher at both sampling timepoints, though not statistically significant (*F*_1,11.2_ = 1.97, *p* = 0.187; Fig. 1D); maximum plant height was greater from the beginning to mid-growth season (precipitation × date: *F*_9, 446_ = 8.18, *p* < 0.001; Fig. 1E), and became indifferentiable between treatments after exponential growth.

### 3.2. Composition of rhizodeposit C consumers

We identified 255 bacterial and 42 fungal consumers of rhizodeposit C, represented by distinct ASVs, across treatments the day after labeling, which rose to 328 bacterial and 47 fungal ASVs four weeks later (Table S5). Detectable rhizodeposit C consumers represented a small fraction of the rhizosphere community, but the percentage of consumers was higher under reduced than average precipitation (bacteria: 8.2% vs. 5.1% and fungi: 2.1% vs. 1.9% of all taxa at the later sampling timepoint). Reduced precipitation increased the number of distinct bacterial consumers by 29% and fungal consumers by 11% the day after labeling, gaps that widened to 68% for bacteria and 13% for fungi four weeks later (Fig. 2A, B; Table S5). Late-exclusive bacterial consumers outnumbered early-exclusive consumers fourfold under reduced precipitation, whereas under average precipitation these numbers were near-equal (Fig. 2A; Table S5). This increase in consumers under reduced precipitation was robust to qSIP threshold choice: applying the more conservative criterion of lower 90% confidence interval > 0 (vs. mean EAF > 0 used throughout) recovered the same pattern of higher consumer richness under reduced precipitation (Fig. S4).

**Fig. 2.**
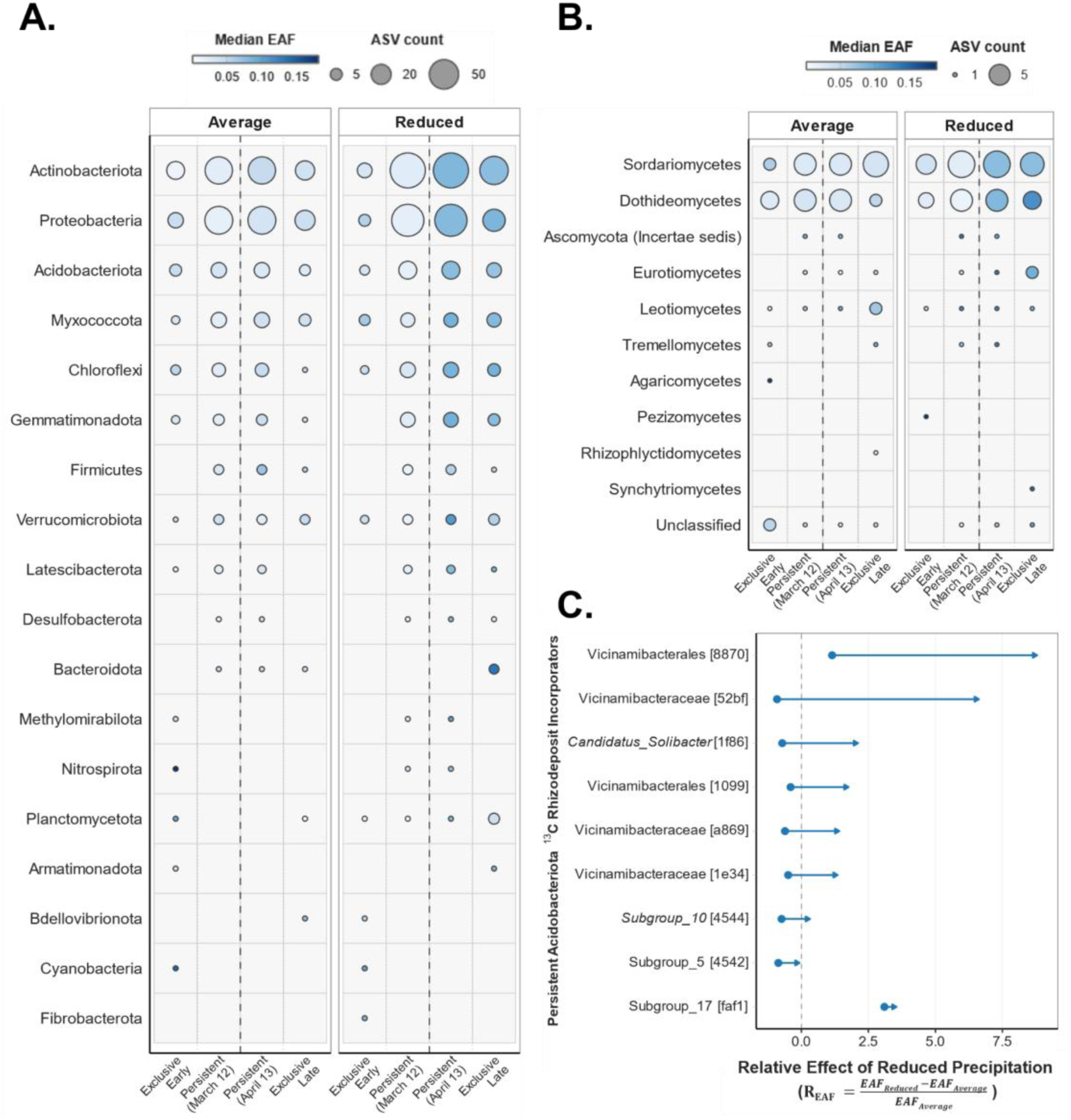
Temporal dynamics of rhizodeposit ¹³C uptake across bacterial and fungal communities under contrasting precipitation treatments in a California annual grassland. Rhizodeposit C was traced via ¹³CO₂ pulse labeling and consumers were identified by qSIP (¹³C incorporation quantified as mean EAF > 0) at two timepoints: immediately after ¹³CO₂ pulse labeling of grasses (March 12) and four weeks later (April 13). Bubble matrices summarize **(A)** bacterial consumers aggregated by phylum and **(B)** fungal consumers aggregated by class, under average and reduced precipitation. Taxa are assigned to one of four temporal categories: exclusive early consumers present only at March 12 (Exclusive Early), persistent consumers present at March 12 (Persistent, March 12), persistent consumers present at April 13 (Persistent, April 13), or exclusive late consumers present only at April 13 (Exclusive Late). Persistent consumers are those that are ^13^C-enriched at both timepoints within a given treatment and therefore appear in both Persistent columns. Bubble size reflects the number of taxa in each phylum– or class–category combination; fill color reflects mean EAF. Ascomycota (Incertae sedis) in panel B refers to confirmed Ascomycota with unresolved class-level placement in the UNITE database; Unclassified refers to taxa with no reliable class-level assignment. Individual taxon identities, full taxonomy, and mean EAF values are provided in Supplementary Data 1. **(C)** Relative effect of reduced precipitation on ¹³C enrichment (R_EAF_) for persistent Acidobacteriota taxa. Each arrow represents a single ASV; the tail marks R_EAF_ at March 12 and the arrowhead marks R_EAF_ at April 13, where R_EAF_ = (EAF_Reduced_ − EAF_Average_) / EAF_Average_. Values to the right of the dashed line (R_EAF_ > 0) indicate higher rhizodeposit C uptake under reduced precipitation. Taxon labels show the lowest resolved taxonomic assignment followed by the first four characters of the taxon identifier in brackets; italic labels indicate genus-level assignments and plain labels indicate family- or order-level assignments. Standard errors for R_EAF_ and full taxonomy are provided in Supplementary Data 2. Results for other persistent consumers are in Figs. S1 and S2.

Bacterial consumers were dominated by Actinobacteriota (134 ASVs) and Proteobacteria (102 ASVs), and fungal consumers were exclusively Ascomycota (51 ASVs). The most ¹³C-enriched bacterial consumer belonged to the genus *Solirubrobacter* (mean EAF = 0.28, indicating that compared with natural abundance, 28% more of C atoms in its DNA were ^13^C, at the later timepoint under reduced precipitation; Supplementary Data 1). The most ¹³C-enriched fungal consumer belonged to the class Sordariomycetes (mean EAF = 0.25 at the later timepoint under reduced precipitation; Supplementary Data 1). Late-exclusive bacterial consumers included Myxococcota (Haliangiales, Myxococcales, Polyangiales) and Bacteroidota (Cytophagales) in both treatments, but were more numerous and more ¹³C-enriched under reduced precipitation (Fig. 2A; Supplementary Data 1). Among fungi, two major fungal classes — Sordariomycetes and Dothideomycetes — dominated rhizodeposit C uptake across both treatments, with higher incorporation under reduced precipitation at the later timepoint (Fig. 2B). A third class, Eurotiomycetes, showed minimal incorporation under average precipitation, but under reduced precipitation, *Penicillium* increased ¹³C enrichment 3.6-fold between timepoints while *Aspergillus* and *Talaromyces* emerged as late-exclusive fungal consumers (Fig. 2B; Supplementary Data 1).

Among the 127 persistent bacterial and fungal consumers (ASVs identified as ¹³C-enriched across both treatments and timepoints), 49% showed higher enrichment under reduced vs. average precipitation at the first timepoint (*R_EAF_* > 0), increasing to 94% after four weeks (95% of bacteria and 100% of fungi), with 80% showing a stronger water-reduction effect over time (Fig. S1, S2). This trend was most pronounced for Acidobacteriota, for which all persistent consumers showed stronger water-reduction effects on enrichment over time (Fig. 2C).

The composition of bacterial consumers, analyzed based on their relative abundances in the total bacterial communities, did not differ significantly between precipitation treatments (PERMANOVA on plot-level centroids: *R²* = 10.3%, *p* = 0.097; Fig. 3A); fungal consumers similarly showed no significant relative abundance response to treatment (*R²* = 10.6%, *p* = 0.089; Fig. 3B). Compositional shifts between timepoints were significant for both groups (within-plot PERMANOVA: bacteria *R²* = 6.5%, *p* = 0.005; fungi *R²* = 3.1%, *p* = 0.032). Relative abundances of the 127 persistent consumers showed no consistent response to water reduction, including for all persistent consumers belonging to Acidobacteriota; the relative effect on abundance (*R_RA_*) was variable and bidirectional between timepoints (Figs. S1, S2).

**Fig. 3.**
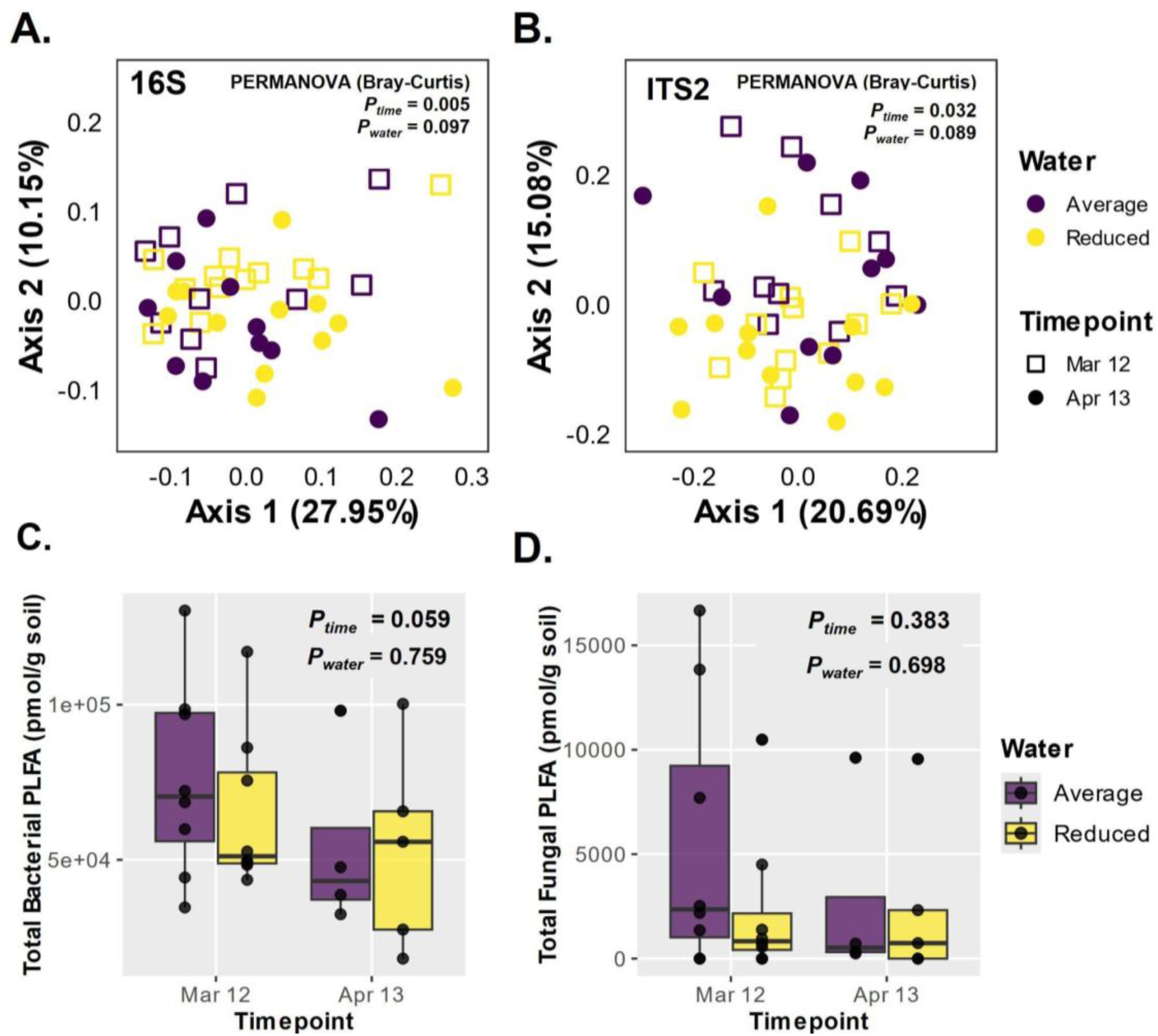
Microbial community composition and biomass under contrasting precipitation treatments in a California annual grassland. PCoA of Bray-Curtis dissimilarities calculated from relative abundances of **(A)** bacterial (16S) and **(B)** fungal (ITS2) rhizodeposit C consumers identified via qSIP (¹³C incorporation quantified as mean EAF > 0) under average (*n* = 10) and reduced (*n* =12) precipitation at two timepoints: immediately after ¹³CO₂ pulse labeling of grasses (March 12 = open squares) and four weeks later (April 13 = filled circles). Total **(C)** bacterial and **(D)** fungal biomass (pmol PLFA g⁻¹ soil) under average (*n* = 8 for March 12 and *n* = 4 for April 13) and reduced (*n* = 8 for March 12 and *n* = 5 for April 13) precipitation. Statistical results are from PERMANOVA on Bray-Curtis dissimilarities with the precipitation effect tested at the plot level (plot centroids) and the timepoint effect tested within plots (panels A and B), and from two-way mixed-effects ANOVA (panels C and D).

Total bacterial and fungal biomass in the rhizosphere, measured by phospholipid fatty acid analysis (PLFA), were unaffected by the precipitation treatment (bacteria: *p* = 0.759; fungi: *p* = 0.698; Fig. 3C, D). Bacterial biomass declined marginally between timepoints (*F*₁,₁₇ = 4.09, *p* = 0.059).

### 3.3. Co-occurrence networks of rhizodeposit C consumers

We constructed ^13^C-informed co-occurrence networks for rhizodeposit C consumers in which each taxon’s relative abundance was weighted by its mean ¹³C enrichment, such that taxon abundance and activity in the consumption of rhizodeposit-derived C both contributed to inferred associations. Networks were larger and more connected at the later timepoint under both precipitation treatments (Fig. 4). The reduced-precipitation network at the later timepoint had at least 37% more nodes and 65% more connections per node (higher average degree) than the other networks, with lower modularity (0.60 vs 0.74–0.77; Table 1; Fig. 4). Bacteria–bacteria associations dominated this network, accounting for 81% of all associations (Table S8). Negative associations were slightly more prevalent relative to the other three networks (38.8% vs 31.8–32.6%; Table 1), with the highest proportions among bacteria (38.5%) and between bacteria and fungi (41.1%). This dense, highly connected reduced-precipitation network at the later timepoint was anchored by a specific cohort of keystone bacteria and fungi. Bacterial broker-hubs — taxa with both high connections and frequent occurrence on the shortest paths linking other taxa in the network — were from Actinobacteriota (unclassified Gaiellales, *67-14*, *Rubrobacter*, and *Conexibacter*) and Proteobacteria (*MND1*) (Fig. 4D, IDs 1-5). Bacterial–fungal associations were dominated by five fungal connectors, taxa that exclusively associated with bacterial partners and not other fungi, from Ascomycota of the genera *Aspergillus*, *Penicillium*, *Pyrenophora*, *Chaetosphaeronema*, and *Trematosphaeria* (Fig. 4D, IDs 6-10).

**Fig. 4.**
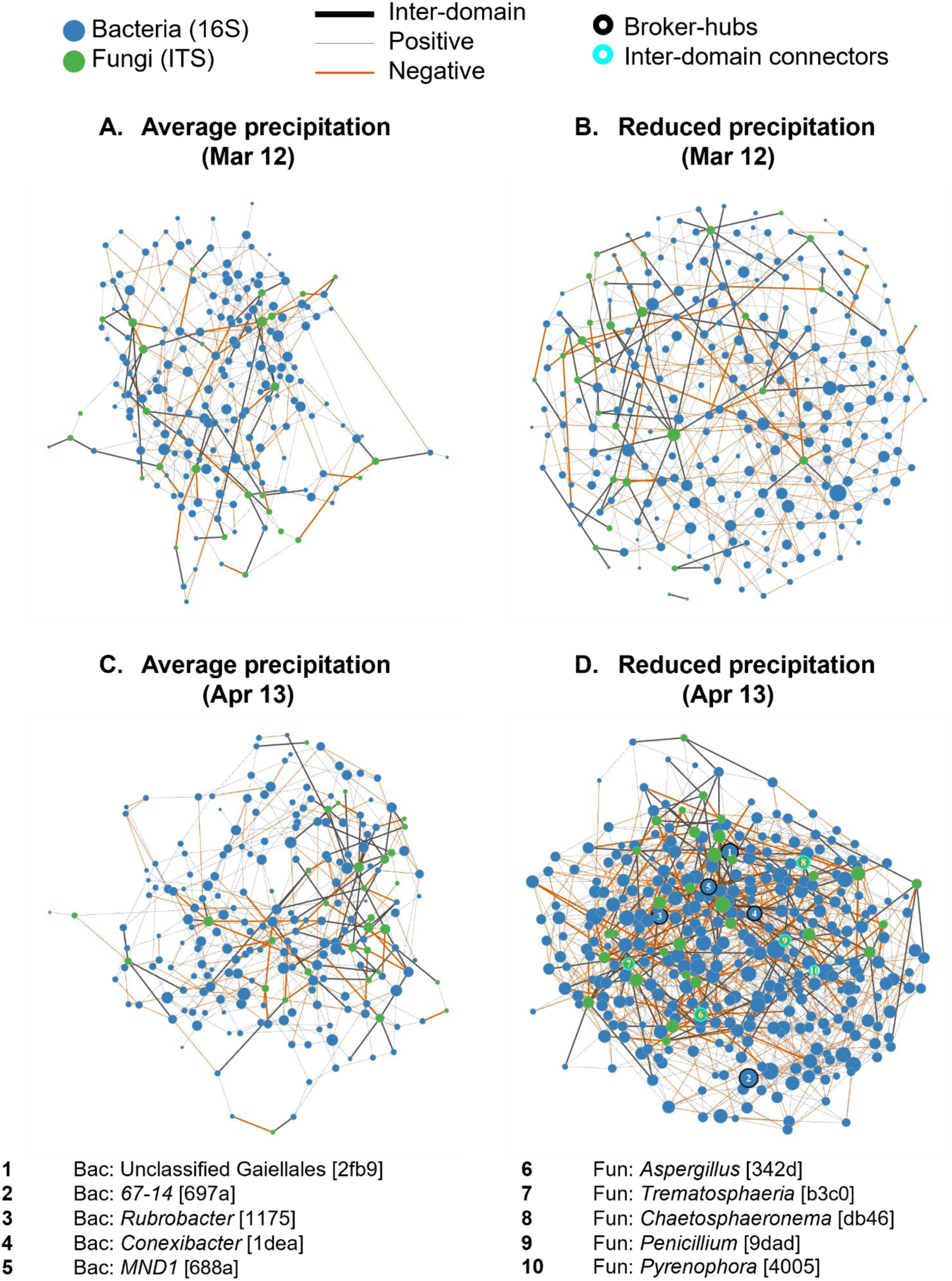
Co-occurrence networks of rhizodeposit C consumers under contrasting precipitation treatments in a California annual grassland at two timepoints. Rhizodeposit C was traced via ¹³CO₂ pulse labeling and consumers identified via qSIP (¹³C incorporation quantified as mean EAF > 0); networks were constructed for rhizodeposit C consumers, with relative abundances weighted by mean EAF to capture activity in rhizodeposit C uptake. Associations were inferred using SpiecEasi for consumers under average (left) and reduced precipitation (right) on March 12 (top) and April 13 (bottom). Nodes represent individual consumers colored by domain: bacteria (blue) and fungi (green); node size is proportional to the number of connections (degree). Edge thickness distinguishes within-domain (thin) from bacterial-fungal (thick) associations; edge color indicates association sign: gray for positive and orange for negative. Node outlines and numbered labels in the April 13 reduced precipitation network highlight keystone roles: black outlines indicate the top five broker-hubs (taxa with both high degree and high betweenness centrality) and cyan outlines indicate the top five bacterial-fungal connectors (taxa where the majority of connections link bacteria to fungi), with betweenness centrality as a tiebreaker. Numbered taxa are listed below the figure at the lowest resolved taxonomic level; brackets contain the first four characters of the taxon identifier for cross-referencing with Supplementary Data 1.

**Table 1.** Global properties of co-occurrence networks for rhizodeposit C consumers under contrasting precipitation treatments in a California annual grassland.

| Precipitation | Timepoint | Taxa (nodes) | Connections (edges) | Negative associations (%) | Connections per node (avg. degree) | Modularity |
| --- | --- | --- | --- | --- | --- | --- |
| Average | March 12 | 200 | 365 | 32.6 | 3.65 | 0.77 |
|  | April 13 | 217 | 433 | 32.3 | 3.99 | 0.75 |
| Reduced | March 12 | 253 | 512 | 31.8 | 4.04 | 0.74 |
|  | April 13 | 346 | 1155 | 38.8 | 6.67 | 0.60 |

For comparison, parallel networks built using only relative abundances of the total community (without ¹³C weighting) recovered a different topology: these total community networks were denser with 3–133% higher average degrees compared to the ^13^C networks, capturing 67.6–95.4% of nodes but only 16.2–33.5% of the edges in the ^13^C networks (Table S11). Total community networks under reduced precipitation showed decreased connectivity (Fig. S3; Table S12), opposite to the pattern observed in ¹³C-informed networks.

### 3.4. Photosynthetic C distribution and modeled solute transport

We tracked the flow of ^13^C across plant and soil pools to test whether its movement into the surrounding soil differed under contrasting precipitation. Atom % ¹³C in shoot biomass was marginally lower under reduced precipitation (*F*_1, 9_ = 4.21, *p* = 0.07) and declined significantly between timepoints under both treatments (*F*_1, 9_ = 30.19, *p* < 0.001; Fig. 5A). Root atom % ¹³C trended, but not significantly, lower under reduced precipitation (precipitation: *F*_1, 13_ = 1.01, *p* = 0.333; Fig. 5B) and higher at the later timepoint (timepoint: *F*_1, 13_ = 2.25, *p* = 0.158). The surrounding soil beyond the root surface, sampled separately from the rhizosphere, contained slightly less ¹³C under reduced precipitation at both timepoints (*F*_1, 9_ = 4.12, *p* = 0.073; Fig. 5C).

**Fig. 5.**
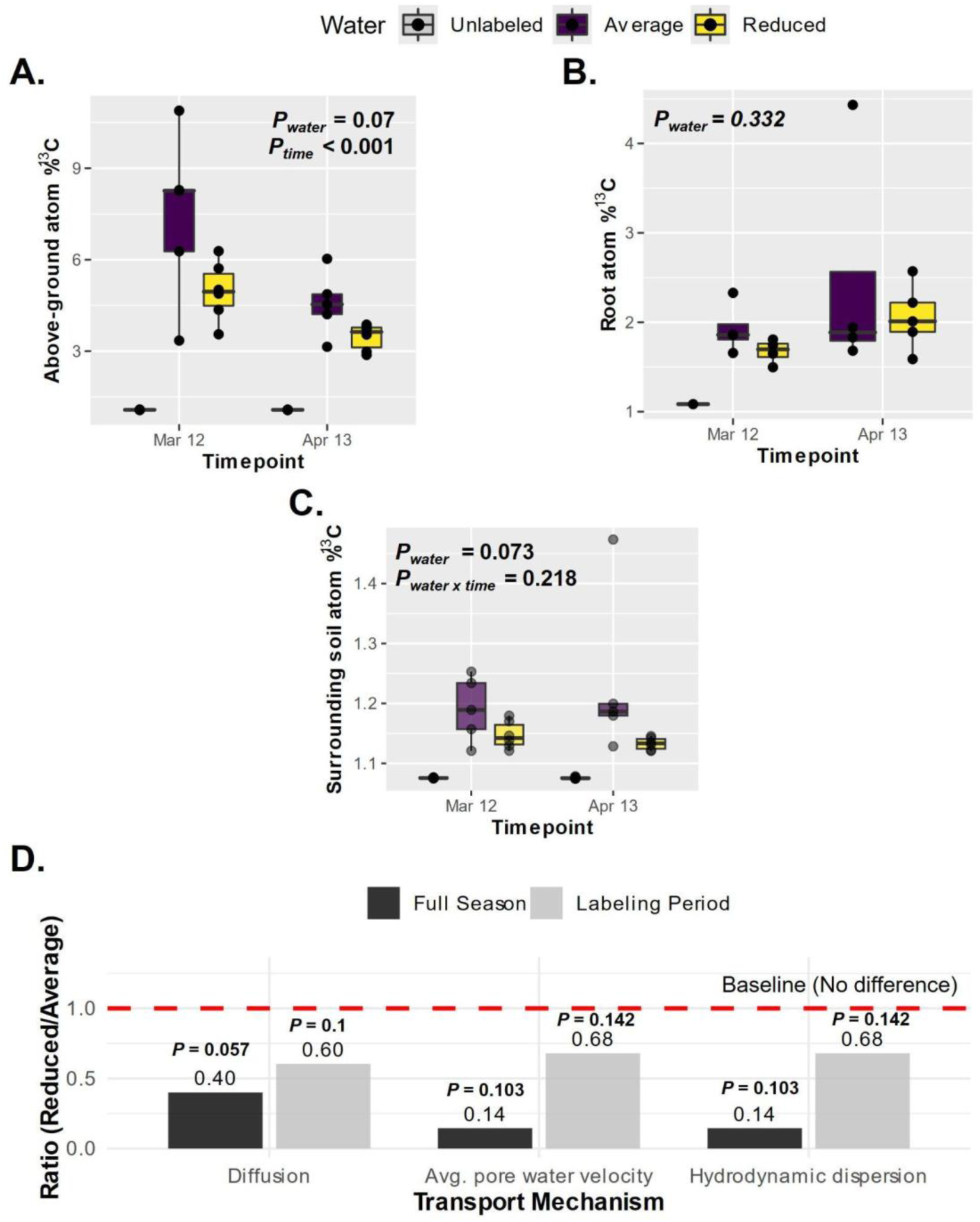
Photosynthetic ¹³C distribution across plant and soil pools and modeled solute transport under contrasting precipitation treatments in a California annual grassland. Plant and soil ¹³C were traced after eight-day ^13^CO_2_ pulse labeling (March 3-11, 2020) during *Avena*’s exponential growth. Atom % ¹³C in **(A)** above-ground biomass (average, purple: *n* = 5; reduced, yellow: *n* = 6), and **(B)** roots (average: *n* = 4; reduced: *n* = 4 at March 12, *n* = 5 at April 13), and **(C)** atom % ¹³C in surrounding soil (average, purple, *n* = 5; reduced, yellow, *n* = 6); unlabeled control samples are shown to illustrate background atom % ¹³C but were excluded from statistical analyses. **(D)** Ratios of modeled transport parameters (reduced/average precipitation) for the full season (black) and the isotope-labeling plus sampling period (gray; March 3-April 13, 2020); values < 1 indicate reduced transport capacity under reduced precipitation. Statistical results are from two-way mixed-effects ANOVA (panels A, B, and C) and Welch’s two-sample t-tests on log-transformed plot-level means (panel D).

To evaluate whether the lower ^13^C in surrounding soil could be due to reduced solute transport, we modeled relative solute transport rates by three dominant physical processes — molecular diffusion, average pore-water velocity, and hydrodynamic dispersion from soil water-retention curves. During the 42-day period between the start of ^13^C labeling and the last sampling, there was 40% lower molecular diffusion (*t*_12.5_ = 1.77, *p* = 0.101; Fig. 5D) and 32% lower average pore-water velocity and hydrodynamic dispersion (*t*_10_ = 1.59, *p* = 0.142) under reduced precipitation. These reductions were even more profound over the entire growing season, with 60% lower diffusion (*t*_10.3_ = 2.14, *p* = 0.057) and 85% lower average pore-water velocity and hydrodynamic dispersion (*t*_13.8_ = 1.75, *p* = 0.103).

## 4. Discussion

### 4.1. Increased number, enrichment, and connectivity of rhizodeposit C consumers under reduced precipitation

Despite substantially lower soil water availability under reduced precipitation, the number of rhizodeposit C consumers, their per-taxon enrichment, and the connectivity among them all increased. The fourfold dominance of late-exclusive over early-exclusive bacterial consumers under reduced precipitation suggests that there were more distinct consumers rather than amplified activities of existing consumers over time with lowered water availability. Several converging processes likely shaped the temporal shifts in the consumer community. From the first to the second sampling timepoint, drier soils could impose stronger physiological constraints on microbial activity. It is likely that the rhizosphere community structure evolved due to shifts in exudation chemistry, as annual grasses moved from exponential growth toward near senescence, a pattern documented in our previous greenhouse work where rhizosphere communities tracked plant developmental stage (Shi et al., 2015; Zhalnina et al., 2018). At the same time rhizodeposit C moved across trophic levels of the soil food web, from primary to secondary consumers, increasing the number of microbes with access to ¹³C. The appearance of bacterial predators (Myxococcota: Haliangiales, Myxococcales, Polyangiales) (Zhang and Lueders, 2017; Petters et al., 2021; C. Wang et al., 2023; Groß et al., 2023) and fungal biomass degraders (Bacteroidota: Cytophagales) (Emmett et al., 2021) among the bacterial consumers exclusively at the later timepoint suggests that rhizodeposit-derived C had moved beyond primary consumers into higher trophic levels of the rhizosphere food web over time. This trophic transfer appears amplified under reduced precipitation.

Root exudation fuels a diverse soil microbial food web, and C fixed during a short pulse label can persist among soil microbial communities through the growing season and potentially longer (Pausch and Kuzyakov, 2018; Fossum et al., 2022). The amplified temporal increase in ¹³C enrichment among persistent consumers under reduced precipitation suggests that these consumers more actively intercepted and incorporated rhizodeposit C near roots. Yet this increase in uptake of rhizodeposit C did not translate into an advantage in population size, indicated by the inconsistent relative abundance change, nor a restructure of the broader consumer communities, indicated by the lack of their compositional shift. The increase in number and enrichment of consumers thus cannot be readily explained by osmotic stress suppression of dominant competitors or altered root exudate chemistry under drying, both of which would be expected to shift the identity of active consumers (Nuccio et al., 2016; Zhalnina et al., 2018) rather than increase their number. Stable microbial biomass despite higher rhizodeposit C uptake under reduced precipitation may reflect two non-exclusive processes: faster microbial turnover with greater replacement of unlabeled by labeled DNA while total biomass remains unchanged, or mineralization of assimilated rhizodeposit C rather than its retention in biomass. The latter is consistent with lower carbon use efficiency (CUE) under drought in the same soil and grass species (Sokol et al., 2024) and lower community growth efficiency (analogous to CUE) under legacy precipitation reduction at this site (Hernandez et al., 2026), though both studies involved more severe water limitation than experienced here. The increased uptake and enrichment of rhizodeposit C consumers indicate that microbial activities were likely not suppressed under reduced precipitation, and argues against reduced decomposition of rhizodeposit C as an explanation for ^13^C accumulation in the rhizosphere. Our measurements did not, however, capture decomposition of unlabeled soil organic matter, which may respond differently to drying.

Reduced precipitation also affected associations among consumers. The ¹³C enrichment-weighted relative abundances quantify taxa activities, linking co-occurrence network structure directly to rhizodeposit C processing. The larger and more connected networks at the later timepoint under both precipitation treatments likely reflect combined effects of strengthened ecological filtering with plant growth (Shi et al., 2015), expanding guilds of active consumers, and trophic transfer of rhizodeposit C over time. The intensified network expansion under reduced precipitation, driven primarily by bacteria–bacteria associations, implies reorganization of rhizodeposit C processing under drier conditions. We posit that taxa occupying central positions in the network, such as bacterial broker-hubs from Actinobacteriota and Proteobacteria, may act as bottlenecks that disproportionately structure associations among consumers and potentially mediate C flow under drier conditions. The positive associations around the five fungal connectors suggest potential facilitative interactions, such as fungal breakdown of complex plant polymers for bacterial use (Yuan et al., 2021). However, the high frequency of negative bacterial–fungal associations indicates that these associations were simultaneously sites of facilitation and competition, likely reflecting both cross-feeding and resource competition for the same rhizodeposit C pool depending on partner identity. Validating the proposed roles of these keystones would require targeted manipulative or imaging-based experiments beyond the scope of this study.

Notably, the positive relationship between rhizodeposit C consumer diversity and network connectivity in this study runs counter to total community networks and to reports of decreased bacterial diversity and network complexity under lower soil moisture (de Vries et al., 2018; Wang et al., 2022). Different mechanisms likely drive these contrasting patterns: total community networks largely reflect environmental filtering imposed by roots (Shi et al., 2016), while ¹³C-informed networks specifically capture associations among taxa actively processing rhizodeposit C. The denser topology but limited edge overlap of total community networks compared with the ¹³C-informed networks suggests that process-relevant associations may be obscured in total community datasets where co-occurrences driven by environmental-filtering dominate. Critically, the increase in negative associations under reduced precipitation was specific to active rhizodeposit C consumers rather than the total community, suggesting competition for the rhizodeposit C pool rather than a general community-wide drought response.

### 4.2. Increased availability of rhizodeposit C near roots: a hypothesis for the increase in microbial rhizodeposit C uptake

The lower ¹³C in shoots and roots under reduced precipitation, though not statistically significant, is consistent with dilution of the fixed ¹³C label across the greater plant biomass observed under this treatment. Because above-ground biomass and plant height were higher under reduced precipitation, a similarly larger root system would spread the same pulse-derived label across more tissue, lowering atom %¹³C in both shoots and roots without requiring reduced C fixation. We therefore interpret the lower shoot and root ¹³C as a consequence of plant size rather than of reduced photosynthetic labeling or belowground allocation.

The lower ¹³C in the surrounding soil (*p* = 0.073) is less easily explained by plant size. One possibility is that less rhizodeposit C reached the surrounding soil under reduced precipitation. However, a larger, actively rhizodepositing root system would be expected to deliver more labeled C outward, not less, making a simple reduction in rhizodeposit supply unlikely. Reduced rhizodeposit input would also imply less labeled substrate available to consumers, which is inconsistent with the increase in consumer ¹³C-rhizodeposit uptake observed under reduced precipitation over time.

A more consistent explanation is that labeled rhizodeposit C remained available to consumers near roots, where it was intensively taken up rather than exported to the surrounding soil. This near-root availability could arise because the plant delivered more rhizodeposit C under reduced precipitation, or because a greater proportion of rhizodeposit C was retained near roots, or both; our data cannot differentiate these. Either process would supply more labeled substrate to consumers at the root surface, accounting for the rise in per-taxon ¹³C enrichment over time.

Our solute transport modeling offers a physical basis for the retention component of this explanation. Rhizodeposit C can be relocated physically via solute transport, and biologically by fungal hyphae or fauna and microbial movement, both of which depend on soil water. The net efflux of rhizodeposits from root to surrounding soil reflects the balance between diffusion outflux, convective and dispersive outflux during precipitation and subsequent drainage and redistribution, and the potential for inward flux driven by root water uptake. The solute transport model, focused on the primary physical dynamic, suggests substantially slower solute transport under reduced precipitation and may explain the reduced ^13^C content in the surrounding soil. As effective diffusion and pore-water velocity are throttled by lower soil moisture, the residence time of rhizodeposits near the root would be extended. This implies confinement of rhizodeposit C near roots and creation of localized hotspots in the rhizosphere (Tecon and Or, 2017; Tecon et al., 2018). Confinement of rhizodeposits near roots has been documented in experimental, imaging, and modeling studies (Carminati et al., 2016; Holz et al., 2018; Zhang et al., 2023; Steiner et al., 2025). Our results suggest this phenomenon may intensify under reduced precipitation in the field.

These transport estimates carry substantial uncertainty given that only three soil cores were used for parameterization, and reflect the *relative* difference in transport potential between treatments rather than absolute fluxes. The model is a simplified physical representation that does not parameterize biochemical sinks such as mineral adsorption or microbial degradation, both of which would further restrict transport *in situ*. The diffusion coefficient used (D₀ = 0.864 cm² day⁻¹) represents a simplified estimate for small organic solutes; larger rhizodeposit compounds would diffuse more slowly, and the model does not account for pore-space occlusion by extracellular polymeric substances in the rhizosphere. Both simplifications likely cause our model to *underestimate* the magnitude of transport restriction, particularly under reduced precipitation. Biological transport pathways were not modeled, but with less connected water films, microbial movement would likely also be restricted. Finally, we did not measure ¹³C in rhizodeposits directly or its distribution with distance from the root, so the concentration of rhizodeposit C near roots remains inferred rather than demonstrated.

Whether through greater delivery or restricted export, the concentration of rhizodeposit C near roots under reduced water could shape the consumer community through spatial isolation. While higher resource availability in the rhizosphere typically promotes competitive exclusion and reduces consumer diversity (Pett-Ridge et al., 2021), fragmentation of soil water films under drier conditions has two parallel consequences that may be relevant here. First, it creates localized hotspots in disconnected microsites where rhizodeposit C concentrates near roots (Tecon and Or, 2017; Tecon et al., 2018); second, it spatially isolates microbial consumers in disconnected microhabitats, weakening competitive exclusion and enabling coexistence among a broader range of taxa (Treves et al., 2003; Tecon and Or, 2017). Both effects intensified as soil water differences between treatments grew over time, consistent with the widening gap in consumer counts between the two sampling times.

This framework also offers an interpretation for the network topology. The slightly elevated frequency of negative associations in this network, particularly among bacteria, potentially reflects competition among consumers sharing the same disconnected microniche for a finite pool of locally concentrated rhizodeposit C. As trophic transfer expanded consumers over time, competition within shared microniches may have intensified. Simultaneously, fragmentation of soil water films may have created more independent disconnected microniches, spatially isolating consumers and reducing between-microniche competition, enabling more diverse taxa to co-exist than would be expected in well-connected soil systems (Treves et al., 2003; Tecon and Or, 2017; Tecon et al., 2018). This is consistent with the observed increase in consumer diversity despite increased negative associations.

The retention of rhizodeposit C in localized hotspots around roots may have implications for grassland SOC stability. At this same site under average precipitation, plant-fixed C was found to associate with the soil mineral fraction within weeks of fixation (Fossum et al., 2022), establishing rhizodeposit C as an important precursor of MAOM. If reduced precipitation limits movement of rhizodeposit C away from roots, as our results suggest, reduced contact with minerals in the surrounding soil may disrupt this rhizodeposit-to-MAOM pathway. This is consistent with evidence that drought reduces MAOM formation from living roots in the same soil and grass species (Sokol et al., 2026), though in a greenhouse setting under more severe water limitation.

### 4.3. Conclusion

By tracing ^13^CO_2_-labeled rhizodeposit C into rhizosphere microbial communities and the surrounding soil, we found that a 50% reduction in precipitation increased microbial uptake of rhizodeposit C without detectable signs of plant stress. Reduced precipitation increased the number of distinct rhizodeposit C consumers and per-taxon ^13^C-enrichment over time during the growing season and intensified associations among consumers in ¹³C-informed co-occurrence networks, even as overall community composition and total biomass remained unchanged. Less ¹³C reached the surrounding soil under reduced precipitation, and modeled solute transport was lower, consistent with rhizodeposit C being retained and intensively processed nearer to roots. These findings suggest that precipitation reduction, even before inducing apparent stress responses in plants, may reshape rhizosphere C cycling through increasing microbial uptake of rhizodeposit C, with potential consequences for rhizosphere-derived contributions to grassland soil C stability (Fig. 6).

**Fig. 6.**
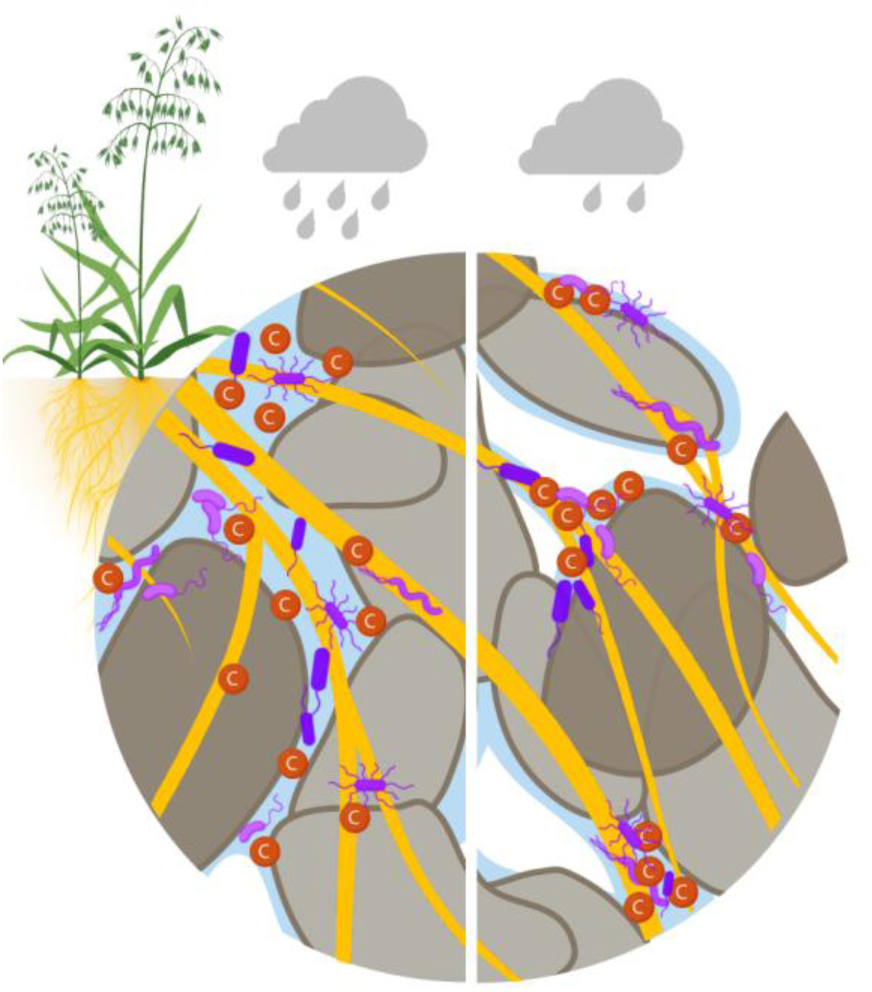
Conceptual diagram illustrating how reduced precipitation may concentrate rhizodeposit C near roots and intensifies its microbial uptake in the grassland rhizosphere. Under average precipitation (left), continuous water films (blue) connect soil pores, facilitating diffusion and transport of rhizodeposit C (red circles) away from roots (yellow) into the soil matrix. Under reduced precipitation (right), fragmentation of water films limits C transport, leading to localized accumulation of rhizodeposit C near roots. We hypothesize that this confinement increases C availability within disconnected rhizosphere microniches, expanding the diversity of microbial consumers and intensifying ecological interactions among rhizosphere microbes (purple).

## Data availability

Raw 16S rRNA gene and ITS2 amplicon sequencing data are deposited in the NCBI Sequence Read Archive (BioProject accession PRJNA1435433). Processed data and intermediate analysis files, including phyloseq objects, qSIP output tables, soil physicochemical data, plant data, PLFA data, and pre-computed co-occurrence network objects, are available on Zenodo (https://doi.org/10.5281/zenodo.18946494). All code used for analyses presented in this manuscript is available on GitHub (https://github.com/nayelazeba/rhizosphere-carbon-precipitation-2020)

## Author contributions

**N.Z.**: Formal analysis, Visualization, Writing – original draft, Writing – review & editing. **K.E.M.**: Methodology, Resources, Investigation, Writing – review & editing. **S.J.**: Data curation, Investigation, Writing – review & editing. **J.Y.**: Methodology, Validation, Writing – review & editing. **S.J.B.**: Data curation, Writing – review & editing. **A.C.**: Data curation. **N.H.N.**: Funding acquisition, Writing – review & editing. **J.Z.**: Funding acquisition, Writing – review & editing. **J.P.R**: Conceptualization, Funding acquisition, Resources, Supervision, Writing – review & editing. **M.K.F.**: Conceptualization, Funding acquisition, Supervision, Resources. **M.Y.**: Data curation, Formal analysis, Writing – original draft, Writing – review & editing.

## Acknowledgements

Our field site exists on territory originally home to the indigenous Pomo Nation. We thank the many individuals who helped in the design, set-up, maintenance, and sampling of this field experiment, including Donald Herman, Carla Yanez, Anne Kakouridis, Evan Starr, Leire Mugica, Nameer Baker, Sarah Baker, Leila Ramanculova, Abel Arellano, Ryan Gini, Max Li, Albert Molina, Angela Hodge, Heejung Cho, Yoni Sher, Ilexis Chu-Jacoby, Lily Law, Caleb Herman, Xiao Chen, Rachel Hestrin, Christian Santos-Medellín, Joanne Emerson, Ella Sieradzki, Alexa Nicolas, Sarick Matzen, Maddie Moore, Cynthia Mancilla, David Sanchez, Tasnim Ahmed, Ricardo Feliciano, Marissa Lafler, Jack Hagen, Natalie Garcia, Albina Khasanova, Erin Nuccio, Kateryna Zhalnina; and the staff at Hopland Research and Extension Center, including Kim Rodrigues, John Bailey, Troy McWilliams, Tom Seward, Zane Kagely, Hannah Bird, Alison Smith, and Shane Feirer. We thank Mike Allen for assistance with density fractionation for qSIP. We thank Hans Singh for his feedback on the manuscript.

## Supplementary Data

**Supplementary Data 1:** Full list of ¹³C-enriched ASVs identified by qSIP across precipitation treatments and timepoints. The sheet provides taxonomy for ¹³C-enriched bacterial and fungal ASVs with mean EAF and CI values at each timepoint, and their category assignment (Early-exclusive, Late-exclusive, or Persistent).

**Supplementary Data 2:** Full list of persistent bacterial rhizodeposit-C consumers (ASVs identified as ^13^C-enriched in both precipitation treatments at both timepoints) with taxonomy and relative effect of reduced precipitation (R_EAF_ and R_RA_) at each timepoint.

**Supplementary Data 3:** Full list of persistent fungal rhizodeposit-C consumers (ASVs identified as ^13^C-enriched in both precipitation treatments at both timepoints) with taxonomy and relative effect of reduced precipitation (R_EAF_ and R_RA_) at each timepoint.

