## Supplementary for "Reduced precipitation alters microbial availability and uptake of grassland rhizosphere carbon"

**Contents**

**1. Supplementary figures S1-S4**

**2. Supplementary tables S1-S12**

**Supplementary figures**

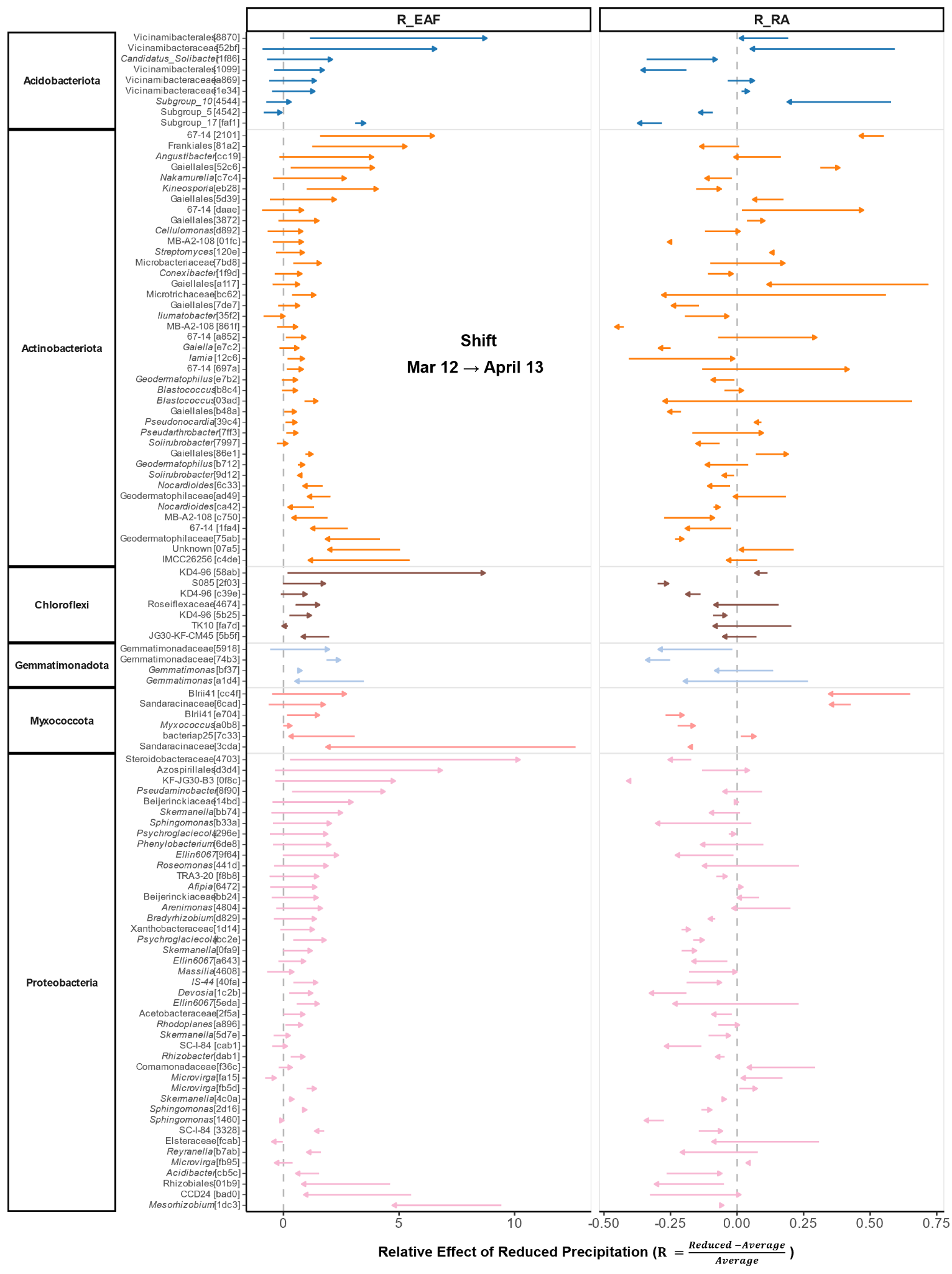

**Fig. S1. Temporal shifts in the relative effect of reduced precipitation on ¹³C enrichment (R_EAF_) and relative abundance (R_RA_) across persistent bacterial consumers of rhizodeposit C.** For each unique bacterial consumer, the arrow tail marks the effect size at the first sampling (March 12) and the arrowhead marks the effect size at the second sampling (April 13). The relative effect R is calculated as (Reduced − Average) / Average, where values to the right of the vertical dashed line (R > 0) indicate higher ¹³C enrichment or relative abundance under reduced compared to average precipitation, and a rightward arrow indicates that the relative effect of precipitation reduction increased between the two samplings. The left panel (R_EAF_) shows the relative effect on ¹³C-rhizodeposit incorporation; the right panel (R_RA_) shows the corresponding relative effect on taxon relative abundance in the total community. ASV labels show the lowest resolved taxonomic assignment followed by the first four characters of the ASV identifier in brackets; italic labels indicate genus-level assignments, plain labels indicate family- or order-level assignments. Standard errors for R_EAF_ and R_RA_ for all persistent ASVs, along with full taxonomy, are provided in Supplementary Data 2.

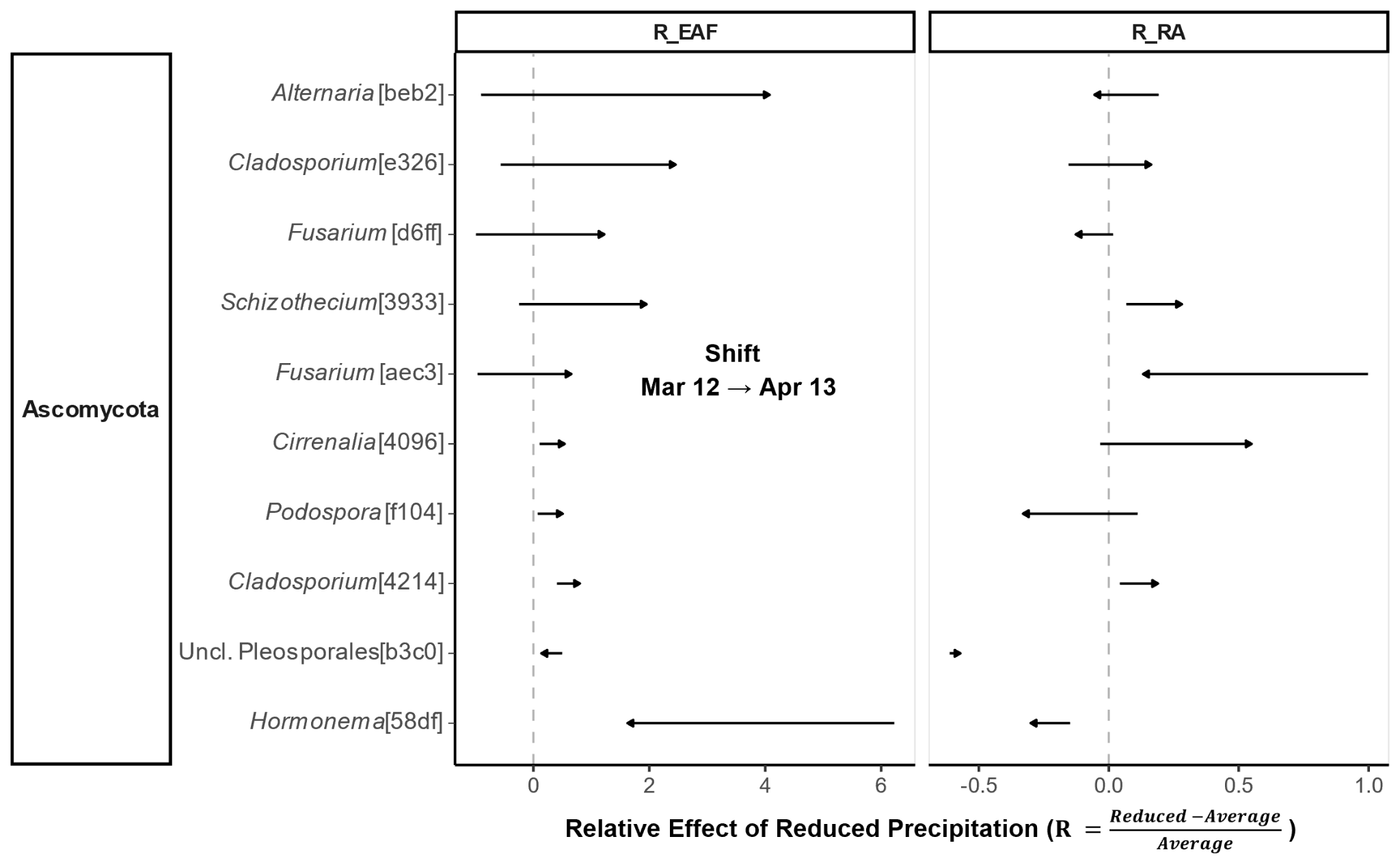

**Fig. S2. Temporal shifts in the relative effect of reduced precipitation on ¹³C enrichment (R_EAF_) and relative abundance (R_RA_) across persistent fungal consumers.** For each fungal consumer, the arrow tail marks the effect size at the first sampling (March 12) and the arrowhead marks the effect size at the second sampling (April 13). The relative effect R is calculated as (Reduced − Average) / Average, where values to the right of the vertical dashed line (R > 0) indicate higher ¹³C enrichment or relative abundance under reduced compared to average precipitation, and a rightward arrow indicates that the relative effect of precipitation reduction increased between the two samplings. The left panel (R_EAF_) shows the relative effect on ¹³C-rhizodeposit incorporation; the right panel (R_RA_) shows the corresponding relative effect on taxon relative abundance in the total community. ASV labels show the lowest resolved taxonomic assignment followed by the first four characters of the ASV identifier in brackets; italic labels indicate genus-level assignments, plain labels indicate family- or order-level assignments. Two ASVs with ambiguous UNITE classifications were assigned taxonomy based on BLASTN searches against the UNITE database. Standard errors for R_EAF_ and R_RA_ for all persistent fungal ASVs, along with full taxonomy, are provided in Supplementary Data 3.

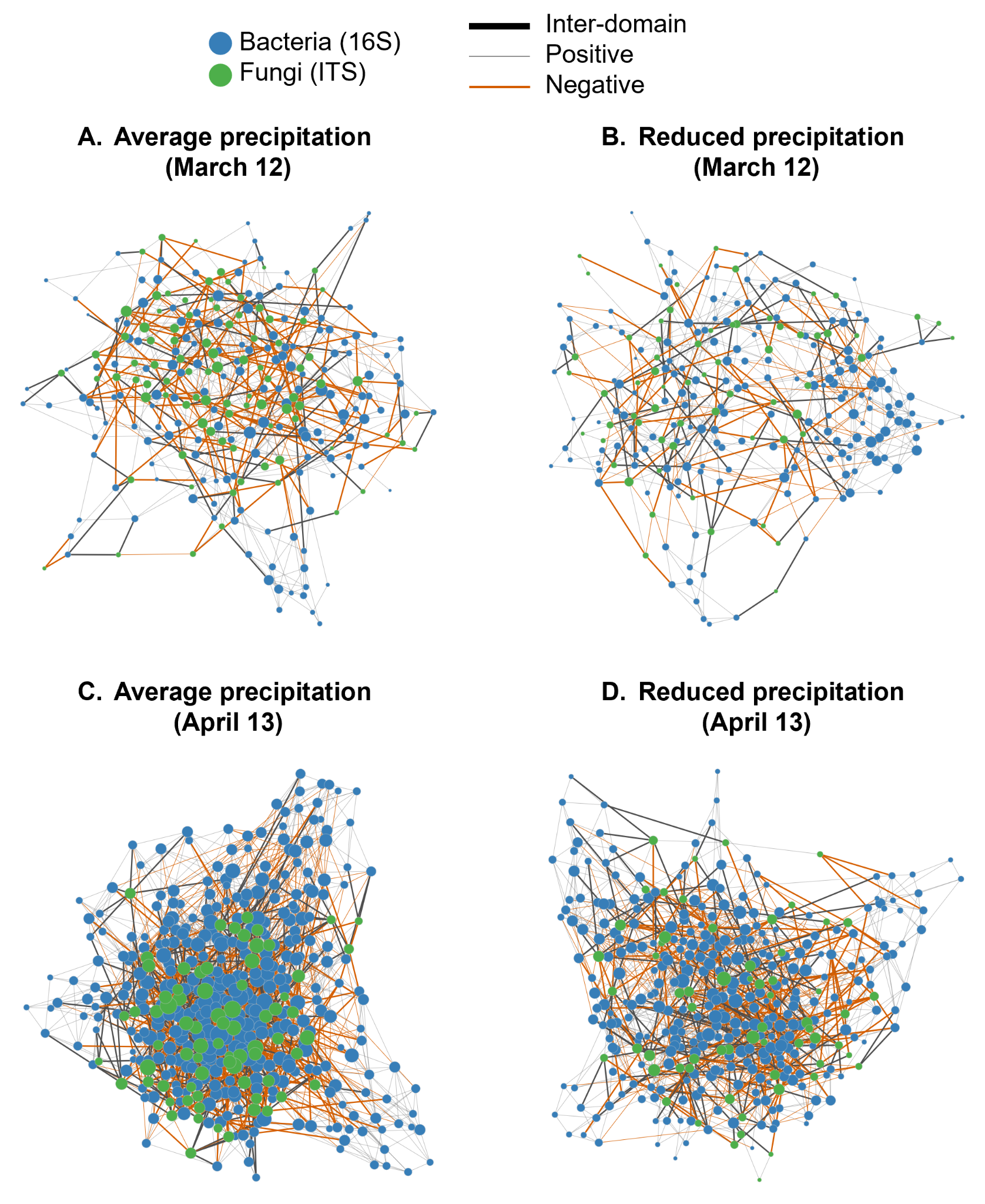

**Fig. S3. Co-occurrence networks of bacterial and fungal taxa across precipitation regimes.** Networks visualize the associations between bacterial and fungal communities under average (left column) and reduced precipitation (right column) on March 12 (top row) and April 13 (bottom row). Networks were inferred using SpiecEasi (MB method) with stability selection. Nodes represent taxa colored by domain: bacteria (blue) and fungi (green). Node size is proportional to degree. Edge thickness distinguishes between intra-domain (thin) and inter-domain (thick) associations. Edge color denotes the sign of the interaction: dark gray for positive and orange for negative.

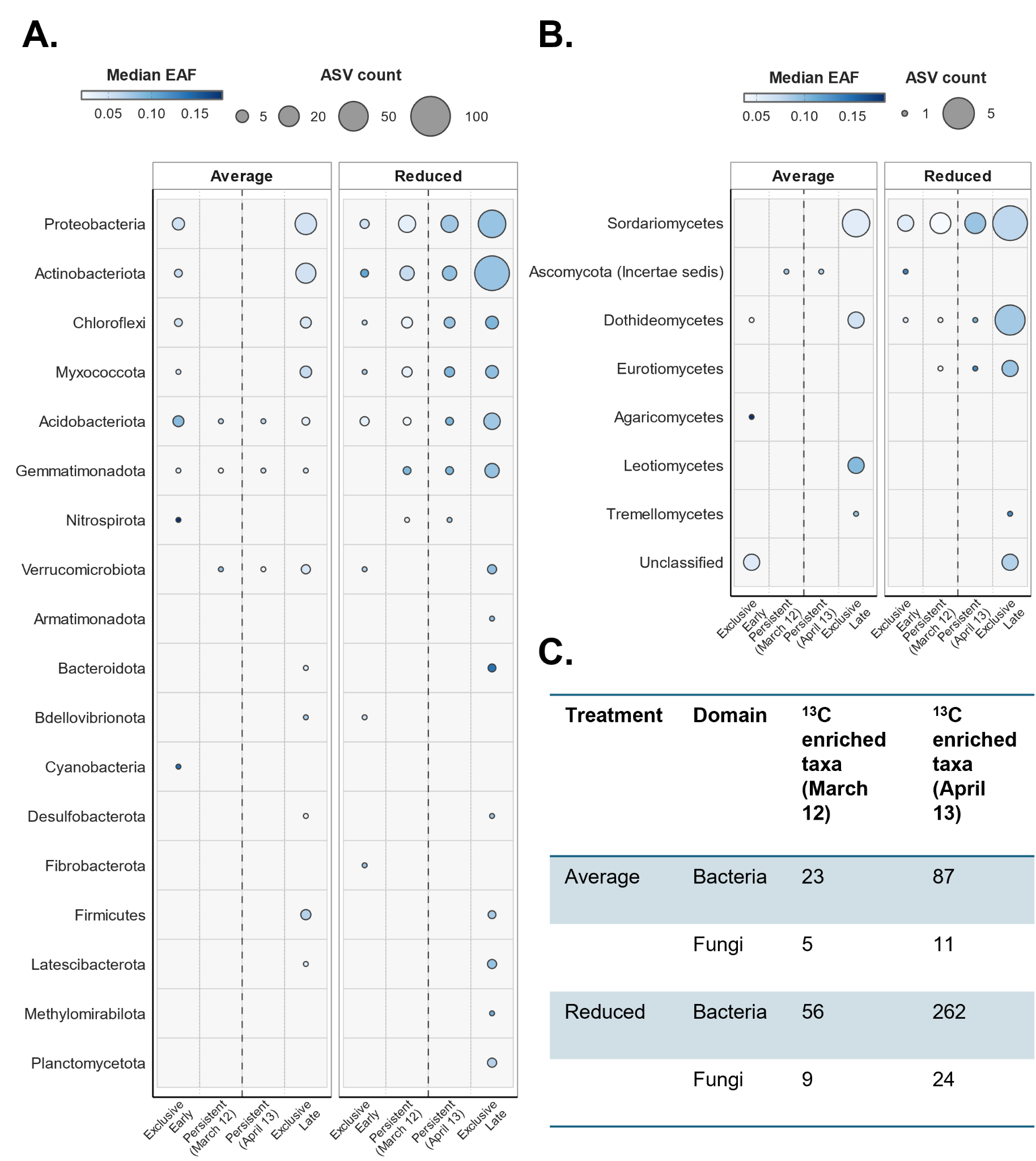

**Fig. S4.** **The temporal patterns of rhizodeposit C incorporation under contrasting precipitation regimes by consumers identified with a more conservative threshold.** Bubble matrices summarizing **(A)** bacterial consumers aggregated by phylum and (B) fungal consumers aggregated by class. Consumers were identified using the conservative criterion in qSIP: lower confidence interval of mean EAF > 0 (lower CI > 0). Temporal categories, color scheme, and bubble scaling follow Fig. 3. **(C)** Total number of bacterial and fungal consumers under average and reduced precipitation at March 12 and April 13, using the lower CI > 0 criterion. The pattern of increased rhizodeposit C consumer diversity under reduced precipitation was consistent with results obtained using mean EAF > 0 (Fig. 3).

**Supplementary tables**

| **Table S1.** Total 16S reads and ASVs recovered from unfractionated DNA | | | | |
| --- | --- | --- | --- | --- |
| Treatment | Timepoint | n | Total reads | ASVs |
| Average | Mar 12 | 10 | 269,990 | 2562 |
|  | Apr 13 | 10 | 516,644 | 3603 |
| Reduced | Mar 12 | 12 | 303,650 | 2411 |
|  | Apr 13 | 12 | 659,683 | 3782 |

| **Table S2.** Total ITS2 reads and ASVs recovered from unfractionated DNA | | | | |
| --- | --- | --- | --- | --- |
| Treatment | Timepoint | n | Total reads | ASVs |
| Average | Mar 12 | 10 | 665,844 | 1552 |
|  | Apr 13 | 10 | 776,129 | 1672 |
| Reduced | Mar 12 | 12 | 797,163 | 1635 |
|  | Apr 13 | 12 | 989,947 | 1729 |

| **Table S3.** Total 16S reads and ASVs recovered from SIP-fractionated DNA | | | | | | | | |
| --- | --- | --- | --- | --- | --- | --- | --- | --- |
| Treatment | Timepoint | n | Fractions | Total reads | ASVs | SIP ASVs | Enriched ASVs (mean EAF > 0) | Enriched ASVs  (lower CI EAF > 0) |
| Average | Mar 12 | 10 | 40 | 1,366,711 | 4518 | 196 | 173 | 23 |
|  | Apr 13 | 10 | 40 | 1,515,856 | 5212 | 200 | 185 | 87 |
| Reduced | Mar 12 | 12 | 48 | 1,688,063 | 4551 | 264 | 223 | 56 |
|  | Apr 13 | 12 | 48 | 1,856,359 | 5306 | 331 | 310 | 262 |

| **Table S4.** Total ITS reads and ASVs recovered from SIP-fractionated DNA | | | | | | | | |
| --- | --- | --- | --- | --- | --- | --- | --- | --- |
| Treatment | Timepoint | n | Fractions | Total reads | ASVs | SIP ASVs | Enriched ASVs (mean EAF > 0) | Enriched ASVs  (lower CI EAF > 0) |
| Average | Mar 12 | 10 | 40 | 2,135,995 | 1539 | 40 | 27 | 5 |
|  | Apr 13 | 10 | 40 | 3,028,600 | 2596 | 37 | 32 | 11 |
| Reduced | Mar 12 | 12 | 48 | 2,397,432 | 1590 | 43 | 30 | 9 |
|  | Apr 13 | 12 | 48 | 3,822,112 | 2758 | 41 | 36 | 24 |

| **Table S5.** Temporal dynamics of ¹³C-enriched ASVs across precipitation treatments | | | | | | | | |
| --- | --- | --- | --- | --- | --- | --- | --- | --- |
| Treatment | Domain | Total | TotalEarly | Total Late | Persistent^a^ | Exclusive Early | Exclusive  Late | Ratio (Ex. Late/Ex. Early) |
| All^b^ | Bacteria | 356 | 255 | 328 | 227 | 28 | 101 | 3.6 |
|  | Fungi | 59 | 42 | 47 | 30 | 12 | 17 | 1.4 |
| Reduced | Bacteria | 339 | 223 | 310 | 194 | 29 | 116 | **4.0** |
|  | Fungi | 46 | 30 | 36 | 20 | 10 | 16 | 1.6 |
| Average | Bacteria | 229 | 173 | 185 | 129 | 44 | 56 | 1.3 |
|  | Fungi | 43 | 27 | 32 | 16 | 11 | 16 | 1.5 |
| ^a^ ASVs enriched at both timepoints within a single treatment. The 127 ASVs persistently enriched across both timepoints and both treatments represent the intersection of these within-treatment persistent pools. | | | | | | | | |
| ^b^ 'All' represents unique ASVs pooled across both precipitation treatments, used to characterize overall temporal dynamics independent of treatment effects. | | | | | | | | |

| **Table S6.** 16S reads and ASVs for ^13^C-network construction after filtering | | | | | | |
| --- | --- | --- | --- | --- | --- | --- |
| Treatment | Timepoint | n | ASVs | % of ASVs in unfractionated DNA | Total reads | % of reads in unfractionated DNA |
| Average | Mar 12 | 10 | 173 | 6.75 | 112,636 | 41.72 |
|  | Apr 13 | 10 | 185 | 5.13 | 200,669 | 38.84 |
| Reduced | Mar 12 | 12 | 223 | 9.25 | 145,616 | 47.96 |
|  | Apr 13 | 12 | 310 | 8.2 | 340,915 | 51.68 |

| **Table S7.** ITS reads and ASVs for ^13^C-network construction after filtering | | | | | | |
| --- | --- | --- | --- | --- | --- | --- |
| Treatment | Timepoint | n | ASVs | % of ASVs in unfractionated DNA | Total reads | % of reads in unfractionated DNA |
| Average | Mar 12 | 10 | 27 | 1.74 | 280,556 | 42.14 |
|  | Apr 13 | 10 | 32 | 1.91 | 377,438 | 48.63 |
| Reduced | Mar 12 | 12 | 30 | 1.83 | 319,650 | 40.1 |
|  | Apr 13 | 12 | 36 | 2.08 | 460,452 | 46.51 |

| **Table S8.** Edge breakdown for the April 13 reduced precipitation ^13^C-network by type and sign | | | | |
| --- | --- | --- | --- | --- |
| Domain category | Association type | Edge count | % of total network | % negative edges |
| Intra-domain | 16S–16S | 936 | 81 | 38.5 |
|  | ITS–ITS | 17 | 1.5 | 29.4 |
| Inter-domain | 16S–ITS | 202 | 17.5 | 41.1 |
| Total |  | 1,155 | 100 | 38.8 |

| **Table S9.** Total 16S reads and ASVs for total community network construction after filtering | | | | | | |
| --- | --- | --- | --- | --- | --- | --- |
| Treatment | Timepoint | n | ASVs | % of ASVs in unfractionated DNA | Total reads | % of reads in unfractionated DNA |
| Average | Mar 12 | 10 | 164 | 6.4 | 114,263 | 42.32 |
|  | Apr 13 | 10 | 357 | 9.91 | 277, 958 | 53.8 |
| Reduced | Mar 12 | 12 | 189 | 7.84 | 148,594 | 48.94 |
|  | Apr 13 | 12 | 303 | 8.01 | 343,922 | 52.13 |

| **Table S10.** Total ITS reads and ASVs for total community network construction after filtering | | | | | | |
| --- | --- | --- | --- | --- | --- | --- |
| Treatment | Timepoint | n | ASVs | % of ASVs in unfractionated DNA | Total reads | % of reads in unfractionated DNA |
| Average | Mar 12 | 10 | 90 | 5.8 | 415,251 | 62.36 |
|  | Apr 13 | 10 | 74 | 4.43 | 452,887 | 58.35 |
| Reduced | Mar 12 | 12 | 64 | 3.91 | 430,015 | 53.94 |
|  | Apr 13 | 12 | 62 | 3.59 | 531,364 | 53.68 |

| **Table S11.** Metrics of node and edge overlap between ^13^C-rhizodeposit and total community networks at each water-treatment × timepoint combination | | | |
| --- | --- | --- | --- |
| Treatment | Timepoint | Node overlap (%) ^a^ | Edge overlap (%) ^b^ |
| Average | Mar 12 | 73.5 | 18.6 |
|  | Apr 13 | 67.6 | 16.2 |
| Reduced | Mar 12 | 95.4 | 33.5 |
|  | Apr 13 | 76.6 | 18.2 |

^a^ Node Overlap %: Calculated as (n_shared_nodes_ / n_13C_nodes_) x 100. This represents the portion of the ^13^C-rhizodeposit community that is also present in the total community network.

^b^ Edge Overlap %: Calculated as (n_shared_edges_ / n_13C_edges_) x 100. This represents the portion of ^13^C-rhizodeposit interactions that are also found in the total community network.

| **Table S12.** Global topological properties of co-occurrence networks constructed from bacterial-fungal communities at each water-treatment × timepoint combination | | | | | | | |
| --- | --- | --- | --- | --- | --- | --- | --- |
| Treatment | Timepoint | Nodes | Total edges | Positive edges (%) | Avg. Degree | Clustering coeff. | Modularity (M) |
| Average | Mar 12 | 254 | 653 | 68.1 | 5.142 | 0.092 | 0.712 |
|  | Apr 13 | 431 | 2009 | 64.5 | 9.323 | 0.109 | 0.561 |
| Reduced | Mar 12 | 253 | 584 | 74.7 | 4.617 | 0.100 | 0.731 |
|  | Apr 13 | 365 | 1257 | 64.3 | 6.888 | 0.095 | 0.628 |
